# Characterizing Cellular Heterogeneity and Transcriptomic Features of Senotype Using Deep Graph Representation Learning

**DOI:** 10.1101/2025.07.01.662364

**Authors:** Anjun Ma, Hao Cheng, Ahmed Ghobashi, Natalia Vanegas-Avendano, Hu Chen, Jhonny Rodriguez-Lopez, Lorena Rosas, Cankun Wang, Jianming Shao, Yi Jiang, Xiaoying Wang, Irfan Rahman, Jose Lugo-Martinez, Dongmei Li, Gloria S. Pryhuber, Ziv Bar-Joseph, Oliver Eickelberg, Melanie Königshoff, Chi Zhang, Dongjun Chung, Rui Chen, Toren Finkel, Ana L. Mora, Mauricio Rojas, Qin Ma

## Abstract

Cellular senescence is a primordial driver of tissue and organ aging, and the accumulation of senescent cells (SnCs) has been implicated in numerous age-related diseases. A major barrier to studying senescence is the rarity and heterogeneity of SnCs, which are not a uniform population but instead comprise diverse senotypes shaped by cell-of-origin and microenvironmental context. Such heterogeneity exceeds what classical senescence hallmarks can resolve at single-cell resolution, motivating the need for computational frameworks that can capture senotype-level diversity intrinsically. Here, we introduce DeepSAS, a deep graph representation learning framework that robustly identifies cell-type-specific SnCs and their senescence-associated genes (SnGs). DeepSAS incorporates a heterogeneous graph that integrates intracellular transcriptional states with intercellular communication cues, enabling the joint inference of senescent cells and senescence-linked genes through attention-based contrastive learning. Applied to public healthy eye and lung atlases, DeepSAS identified SnCs whose proportions positively correlate with aging. From in-house idiopathic pulmonary fibrosis (IPF) patient scRNA-seq data, DeepSAS detected 1,678 SnCs (out of 24,125 cells) and 263 SnGs across 26 cell types, including 43 SnGs that are uniquely associated with a single cell type. We generated high-resolution Xenium spatial transcriptomics data to further validate SnGs in IPF, revealing *NFE2L2* as a SnG specifically enriched in *CTHRC1*+ fibroblasts. Notably, the *ex vivo* bleomycin-induced senescence in human precision-cut lung slice (hPCLS) samples similarly identified *NFE2L2* as an SnG in *CTHRC1*+ fibroblasts, albeit with stronger transcriptional signals, suggesting mechanistic differences in senescence cells associated with chronic and acute injury. Overall, DeepSAS uncovers distinct senescence programs and infers cell-type-specific SnGs that are difficult to resolve using existing marker-based approaches. We believe it offers a generalizable and translationally relevant strategy for advancing senescence biology and therapeutic development.

## INTRODUCTION

Cellular senescence is recognized as one of the main hallmarks of aging and a key contributor to a wide spectrum of age-related diseases, including cancer, atherosclerosis, Alzheimer’s disease, and idiopathic pulmonary fibrosis (IPF)^1,2^. Senescent cells (SnCs) are classically defined by stable cell-cycle arrest and a distinct senescent-associated secretory phenotype (SASP)^3,4^, and their accumulation in tissues has been linked to functional decline and disease progression^5^. Despite their broad relevance, how senescence is encoded across different cell types and biological contexts remains incompletely understood^6^. Historically, senescence has been studied primarily at the bulk tissue level, implicitly treating SnCs as a uniform cellular phenotype. The advent of scRNA-seq enabled finer resolution; yet most single-cell studies continue to model senescence as a homogeneous cell state defined by elevated expression of canonical markers such as *CDKN1A* (*i.e.*, p21) and *CDKN2A* (*i.e.*, p16)^7^. This marker-centric view oversimplifies senescence biology and falls short of capturing its intrinsic heterogeneity across cell types, tissues, and disease states. In reality, every SnC originates from a normal cell and retains transcriptional features of its lineage and microenvironmental context^8^. Although SnCs may share hallmark features such as G1 arrest, lysosomal accumulation and activation, and SASP expression, their underlying senescence programs are shaped by cell-intrinsic transcriptional states and intercellular cues. This layered heterogeneity gives rise to diverse senotypes, representing distinct combinations of senescence triggers, molecular programs, and functional properties embedded within individual cell types and not adequately resolved by existing approaches^9–11^. Curated senescence hallmark gene sets capture only a fraction of senescence-associated variation at single-cell resolution and often are insufficient to identify cell-type-specific senescence programs^6,12^.

Accurate characterization of senotypes at single-cell resolution, therefore, requires computational frameworks that can identify SnCs across diverse cell types, together with their cell-type-specific senescence-associated genes (SnGs), in an unbiased and scalable manner. This task remains challenging for several reasons. SnCs are rare and heterogeneous, making it difficult to distinguish them from non-senescent cells within the same lineage. Moreover, many SnCs exhibit reduced global transcriptional activity^13^, increasing the risk of misclassification as low-quality cells and limiting the detection of subtle, cell-type-specific SnG^14^. Existing computational methods often rely on predefined senescence hallmarks derived from bulk data or focus on specific tissues or diseases, limiting their generalizability^9–11^. Many existing methods apply global scoring schemes that assume a uniform senescence phenotype across cell types; other approaches adopt sequential workflows that first identify senescent cells and then perform differential expression analysis, a strategy that introduces sensitivity to thresholding and reduces power to detect lowly expressed senescence-associated genes^14,15^.

To address the above challenges, we developed DeepSAS, a deep graph representation learning framework for identifying SnCs and SnGs that resolves senescence at single-cell resolution while preserving cell-type specificity. DeepSAS is built on the principle that senescence is embedded within cell-type-specific transcriptional contexts rather than manifesting as a global cellular state. By constructing a heterogeneous graph that integrates intracellular transcriptional states with intercellular cell–cell communication (CCC) cues, DeepSAS enables joint inference of SnCs and SnGs within a unified learning framework. Through attention-based graph modeling and contrastive learning, DeepSAS captures subtle yet biologically meaningful differences between senescent and non-senescent cells within the same cell type, without requiring predefined senescence markers or phenotype labels. This design enables DeepSAS to robustly characterize heterogeneity in senescence across cell types, tissues, and disease contexts, providing a foundation for systematic discovery of cell-type-specific senescence programs.

## RESULTS

### Design Principles and Evaluation of DeepSAS

Unlike prevailing models that conceptualize SnCs either as a homogeneous population or as a distinct cell state separated from lineage identity, DeepSAS explicitly models SnCs as heterogeneous subpopulations that retain the transcriptional and functional characteristics of their cell of origin (**Fig. 1a**). This formulation leads to a fundamental modeling hypothesis: within a given cell type, SnCs are more similar to each other than to SnCs from other lineages, while remaining distinguishable from non-senescent counterparts through subtle but structured transcriptional programs. Accordingly, DeepSAS assumes that SnCs within the same cell type exhibit smaller embedding distances relative to non-SnCs from the same lineage (d_1_<d_3_ in **Fig. 1b**) and to SnCs from different lineages (d_1_<d_2_ in **Fig. 1b**), enabling simultaneous resolution of SnCs identity and cell-type specificity. The framework has the following four unique features (**Online Methods**).

**Fig. 1.**
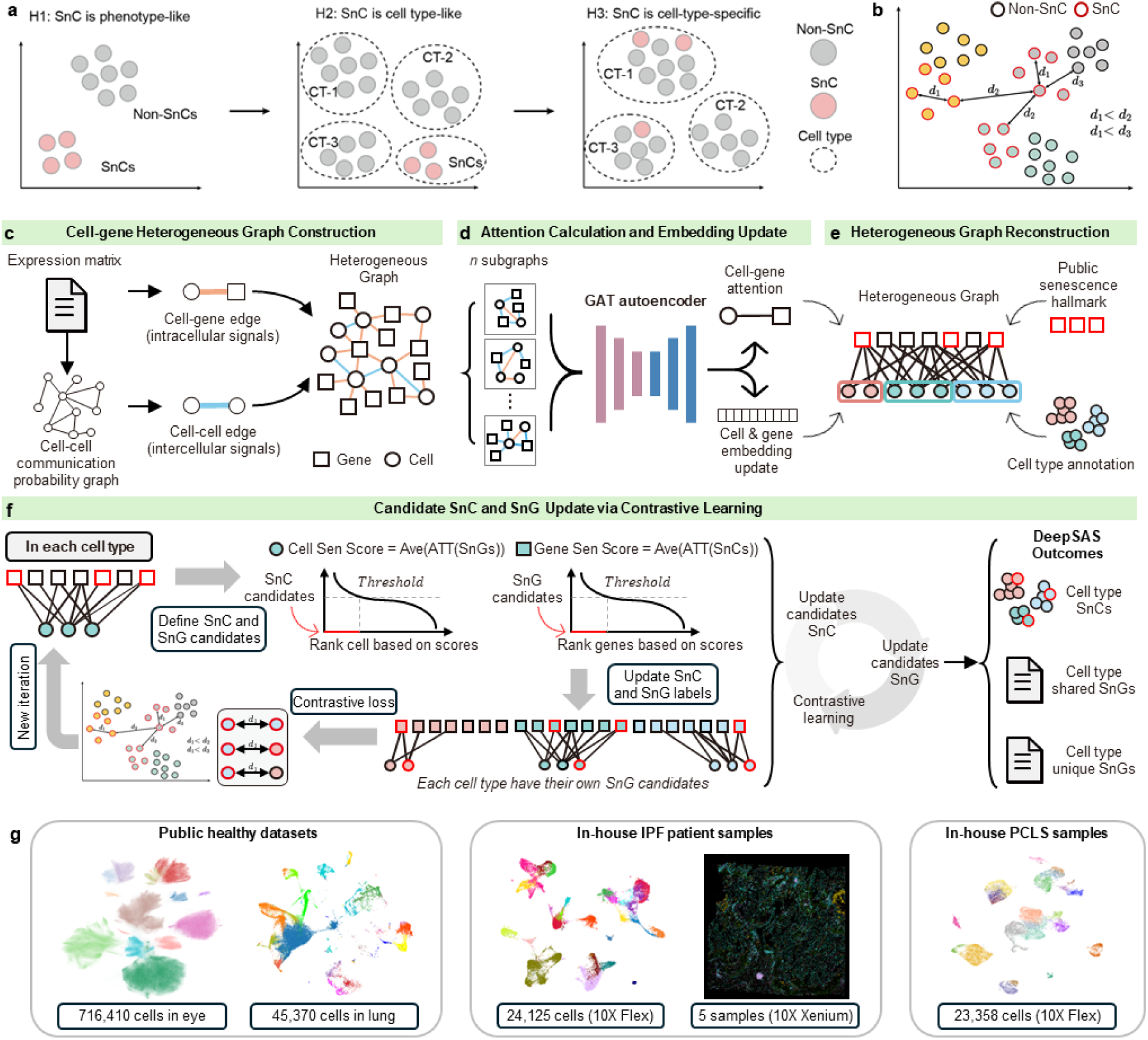
Hypothesis overview and workflow design of DeepSAS. **(a)** Conceptual models of senescent cell identity: Hypothesis 1 (H1) treats SnCs as a uniform phenotype; Hypothesis 2 (H2) models SnCs as a distinct cell type; Hypothesis 1 (H3; our hypothesis) considers SnCs to be cell-type-specific subpopulations requiring context-aware identification. **(b)** DeepSAS assumes SnCs within the same cell type are more transcriptionally similar to each other than to SnCs from other lineages, enabling within-type separation of SnCs and non-SnCs. **(c)** Construction of a heterogeneous cell–gene graph integrating intracellular and intercellular information. Cells and genes are modeled as distinct node types. Cell–gene edges represent intracellular transcriptional signals derived from gene expression, whereas cell–cell edges encode intercellular communication probabilities inferred from ligand–receptor interactions. This unified graph structure enables simultaneous inference of SnCs and SnGs without imposing a predefined identification order. **(d)** Graph attention–based autoencoder for joint embedding of cells and genes. A graph attention network updates latent embeddings for both node types while learning attention weights that capture context-dependent cell–gene associations and cell–cell relationships. **(e)** Heterogeneous graph reconstruction guided by external senescence knowledge. Public senescence hallmark gene sets are incorporated as weak guidance to constrain the search space for senescence-associated programs within each cell type. Attention-derived relationships are used to reweight graph connections and compute senescence scores for both cells and genes. **(f)** An iterative contrastive learning framework operating within individual cell types. SnC and SnG candidates are iteratively updated based on attention-derived senescence scores and contrastive loss, which enforces separation between SnCs and non-SnCs while preserving cell-type structure. This self-supervised process continues until convergence. Outputs of DeepSAS. The framework produces cell-type-specific SnC assignments, cell-type-shared and cell-type-unique SnGs, and continuous senescence scores at both the cellular and gene levels. **(g)** Overview of datasets and application scenarios used to evaluate DeepSAS, including large-scale public healthy human eye and lung atlases, in-house IPF patient scRNA-seq data, and hPCLS scRNA-seq data.

First, DeepSAS constructs a heterogeneous graph that integrates intracellular transcriptional signals with intercellular CCCs into a unified representation. This design is motivated by the observation that senescence is not solely defined by cell-intrinsic transcriptional changes but is reinforced and modulated by microenvironmental cues mediated through SASP^3,4^. Intercellular communication shapes how senescent states are stabilized, propagated, or constrained within specific cellular niches, thereby reinforcing the cell-type specificity of senescence programs^16^. To explicitly encode both cell-intrinsic transcriptional states and extrinsic microenvironmental influences in a unified representation, cells and genes are modeled as distinct node types, connected through weighted cell–gene edges derived from expression profiles and cell–cell edges inferred from SASP-related CCC probabilities (**Fig. 1c**). Importantly, this cell–gene heterogeneous formulation allows SnCs and SnGs to be inferred jointly, rather than through sequential steps that impose artificial dependencies between cell identification and gene discovery.

Second, DeepSAS employs a graph attention transformer (GAT)–based autoencoder to update latent embeddings for both cells and genes. Meanwhile, the GAT also learns cell-gene attentions which quantify neighborhood-dependent importance based on local graph topology and embedding similarities (**Fig. 1d**). The use of graph attention enables the model to adaptively weight gene contributions and cell–cell interactions in a cell-type-aware manner. These learned attention scores form the basis for downstream senescence scoring, providing a representation that is more robust to transcriptional sparsity and noise than activity scores computed solely from gene expression levels.

Third, DeepSAS reconstructs a weighted cell-gene bipartite graph by incorporating external knowledge, including curated public senescence hallmark gene sets and cell-cycle markers, as supervisory signals (**Fig. 1e**). Rather than serving as fixed labels, these external gene sets guide the model toward biologically plausible regions of the feature space, effectively constraining the search for senescence-associated programs within each cell type. This strategy allows DeepSAS to leverage prior biological knowledge while retaining the flexibility to discover novel, cell-type-specific SnGs that are not captured by existing hallmarks.

Finally, DeepSAS adopts an iterative contrastive learning framework that refine SnC and SnG candidates in each cell type (**Fig. 1f**). In each iteration, SnC candidates are selected based on attention-derived senescence scores, and SnG candidates are updated accordingly. A contrastive loss function enforces separation between SnCs and non-SnCs within the same lineage while preserving cell-type structure. This iterative process continues until convergence, enabling cell-type-specific senescence programs to emerge in a self-supervised manner. The final outputs of DeepSAS include cell-level senescence labels and continuous senescence scores, as well as cell-type-specific SnGs that distinguish SnCs and non-SnCs within the same cell type.

To evaluate the robustness and performance of DeepSAS, we conducted systematic benchmarking analyses, including ablation studies, subsampling tests, and comparisons with existing senescence identification approaches. Ablation studies (**Supplementary Fig. S1**) examined the impact of three major design choices: the number of genes used to construct the heterogeneous graph, the inclusion of CCC information, and the use of different hallmark gene lists for labeling initial SnG candidates. The results showed that incorporating CCC edges and integrating multiple hallmark gene sets improved model performance, while varying the number of genes had minimal impact. Robustness tests using different subsampling manners further demonstrated that DeepSAS maintains relatively consistent outputs against phenotype influences, confirming its resilience to sample imbalance and noise (**Supplementary Fig. S2** and **Table S1**). Due to the absence of an established gold standard for single-cell senescence prediction and the lack of existing tools capable of inferring cell-type-specific SnGs (see detailed justification in the **Online Methods**), we focused our comparative evaluation on SnC detection across tools applied to the same datasets (**Supplementary Fig. S3** and **Table S2**). DeepSAS produced biologically plausible predictions of SnC across cell types (average SnC proportion is 7.04%, S^2^=3.36). In contrast, other methods often returned unrealistic results, either predicting an excessively large number of SnCs in multiple cell types (e.g., average SnC proportion is 56.26%, S^2^=1071.72) or generating globally low SnC proportions with limited statistical power (e.g., average SnC proportion is 0.47%, S^2^=0.69). These comparisons reinforce the reliability of DeepSAS in capturing rare but meaningful senescent subpopulations across diverse cellular contexts. We later applied DeepSAS across multiple public and in-house datasets, including large-scale healthy human eye and lung atlases, IPF patient scRNA-seq data, and human precision-cut lung slice (hPCLS), demonstrating its broad applicability across tissues, diseases, and experimental platforms (**Fig. 1g**).

### DeepSAS Reveals Cell-Type-Specific Senescence Patterns Along with Aging in Healthy Tissues

We first applied DeepSAS to a comprehensive human retinal pigment epithelium (RPE) and choroid single-cell atlas comprising 716,410 cells collected from 53 healthy human donors, spanning from infancy to 93 years of age (**Fig. 2a** and **Supplementary Fig. S4; unpublished data, see Data Availability**). This dataset provides an ideal testbed to evaluate how senescence inferred by DeepSAS relates to aging across diverse cell types in the absence of overt disease. To enable scalable analysis at this resolution, the dataset was randomly partitioned into 29 balanced batches while preserving cell-type proportions across batches. Across the atlas, DeepSAS identified 35,185 SnCs, representing 4.91% of all cells (**Fig. 2b**). SnCs were detected across all major cell types, with fibroblasts, RPE cells, and endothelial cells exhibiting the highest SnC proportions. Importantly, SnC proportions were consistent across batches, indicating that DeepSAS predictions were robust to batch partitioning and not driven by technical variation (**Fig. 2c** and **Supplementary Fig. S5**). We next examined the relationship between SnC abundance and donor age. Adult donors exhibited a significantly higher proportion of SnCs compared with pediatric donors (**Fig. 2d** and **Supplementary Fig. S6**). Within fibroblast, RPE, and endothelial populations, SnC proportions increased progressively with donor age, revealing a positive correlation between cellular senescence and aging at the cell-type level (**Fig. 2e**). These observations support the biological relevance of DeepSAS predictions and are consistent with the concept that aging-associated cellular stress leads to the gradual accumulation of SnCs within susceptible cell types. In parallel, DeepSAS inferred cell-type-specific SnGs within this RPE and choroid atlas (**Supplementary Tables S3-S4**). These SnGs were embedded within the transcriptional context of individual cell types and included both shared and cell-type-specific signatures, providing a resource for future mechanistic investigation of senescence in aging-related retinal disorders such as age-related macular degeneration, cataracts, and glaucoma.

**Fig. 2.**
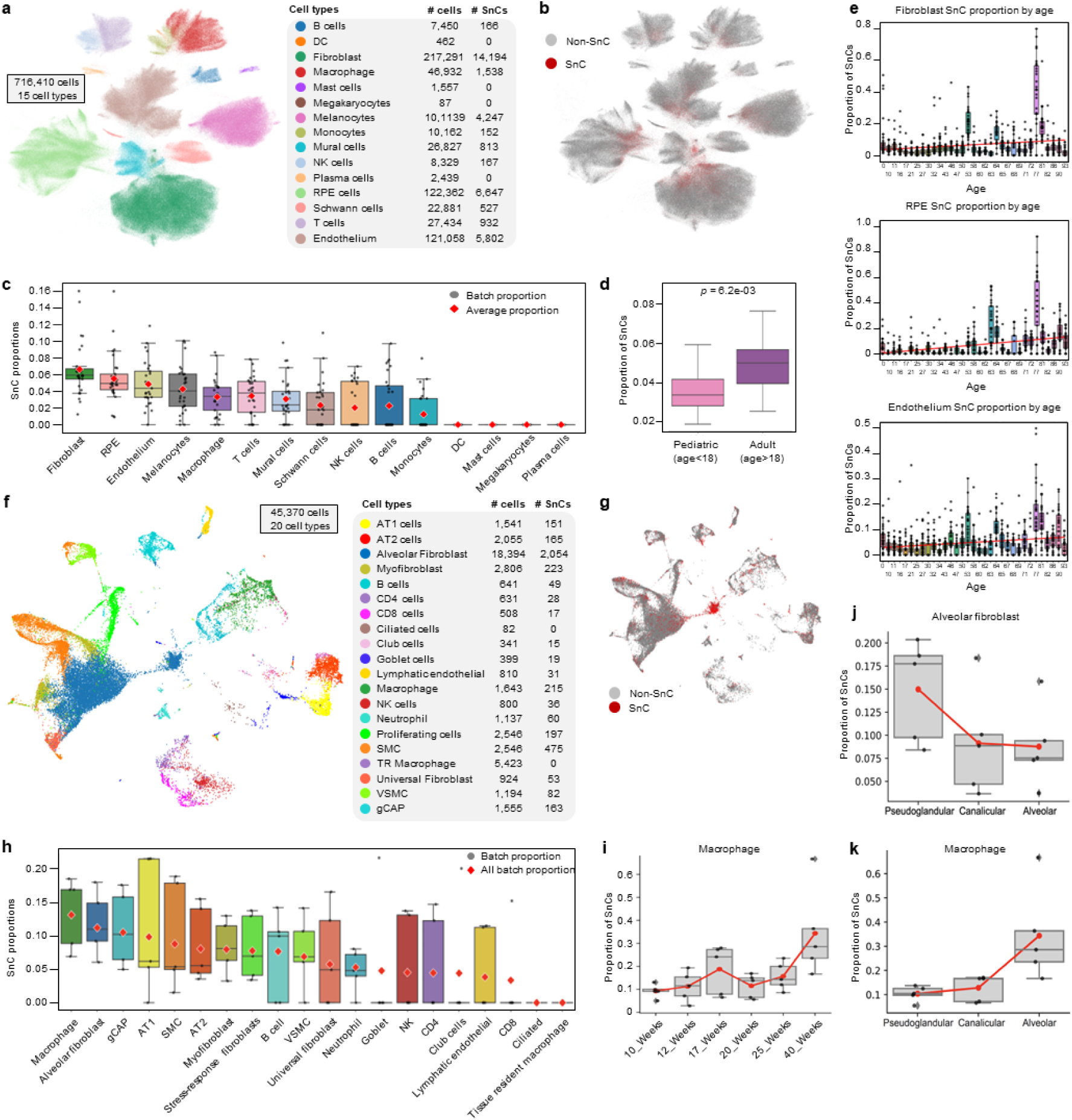
DeepSAS identifies cell-type-specific SnCs associated with aging in healthy human tissues. **(a)** UMAP of the healthy human RPE and choroid single-cell atlas, comprising 716,410 cells annotated into 15 cell types. **(b)** UMAP of the healthy human RPE and choroid single-cell atlas, colored by SnC (red) and non-SnC (grey). **(c)** Comparison of SnC proportions between pediatric and adult donors across all cells in the RPE and choroid atlas. **(d)** SnC proportion as a function of donor age in representative RPE and choroid cell types. **(e)** Cell-type-specific enrichment of SnCs in adult compared with pediatric donors, demonstrating aging-associated senescence within individual lineages. **(f)** UMAP of public healthy lung scRNA-seq dataset comprising 45,370 cells annotated into 20 cell types. **(g)** UMAP of public healthy lung cells, colored by SnC (red) and non-SnC (grey). **(h)** Proportion comparison of SnCs in each of the cell types. **(i)** Changes of SnC proportions in macrophages across different gestational weeks. **(j-k)** Changes of SnC proportions in Alveolar fibroblast and Macrophages across lung developmental stages, representatively.

To assess whether the SnC patterns identified by DeepSAS generalize across tissues while accounting for distinct biological contexts, we applied it to a public healthy human fetal lung scRNA-seq dataset (**Fig. 2f**), comprising 45,370 cells across 20 annotated cell types^17^. In contrast to the retinal analysis, where SnC proportions increased with donor age, fetal lung development represents a setting in which senescence serves a programmed, stage-specific role during organogenesis through the process of developmental senescence^18,19^. Accordingly, we analyzed fetal samples spanning different gestational weeks, which were manually consolidated into the major prenatal lung developmental stages, pseudoglandular, canalicular, and alveolar (**Supplementary Fig. S7**). DeepSAS identified SnCs across multiple lung cell types, including macrophages, alveolar fibroblasts, and smooth muscle cells (**Fig. 2g-h**). Notably, alveolar fibroblasts exhibited higher SnC proportions during the pseudoglandular stage that decreased toward the alveolar stage (**Fig. 2j**), consistent with the transient and developmentally regulated nature of mesenchymal senescence during lung morphogenesis^20,21^. In contrast, macrophages displayed a progressive increase in SnC proportions across advancing developmental stages (**Fig. 2i-k**), indicating that senescence dynamics can differ substantially across cell types even within the same tissue. Together, these results indicate that DeepSAS captures biologically coherent, cell-type-specific senescence programs whose temporal trajectories reflect the underlying biological context, aging-associated accumulation in adult tissues versus stage-restricted deployment during development, rather than enforcing a uniform directionality of senescence across systems.

### DeepSAS Uncovers Functional Heterogeneity and Signaling Dynamics of SnCs In IPF Lungs

We next applied DeepSAS to an in-house 10X Flex scRNA-seq dataset from human IPF lungs, comprising 24,125 cells across 26 annotated cell types from six donors (**Fig. 3a** and **Supplementary Table S5**). This dataset provides a stringent disease-context benchmark to evaluate whether DeepSAS can resolve senescence heterogeneity across diverse cellular compartments without using sample or phenotype information during model training. DeepSAS identified 1,678 SnCs, accounting for 6.96% of all cells (**Fig. 3b**), where SnCs were distributed across epithelial, immune, mesenchymal, and endothelial compartments, rather than confined to a single lineage. When stratified by sample origin, the vast majority of SnCs were derived from IPF samples, with a small fraction observed in healthy aged samples and virtually none detected in healthy young samples (**Fig. 3c** and **Supplementary Table S6**). This distribution is consistent with established biological expectations that cellular senescence is enriched in fibrotic lungs and increases with age, supporting the specificity of DeepSAS predictions in a disease setting^22,23^.

**Fig. 3.**
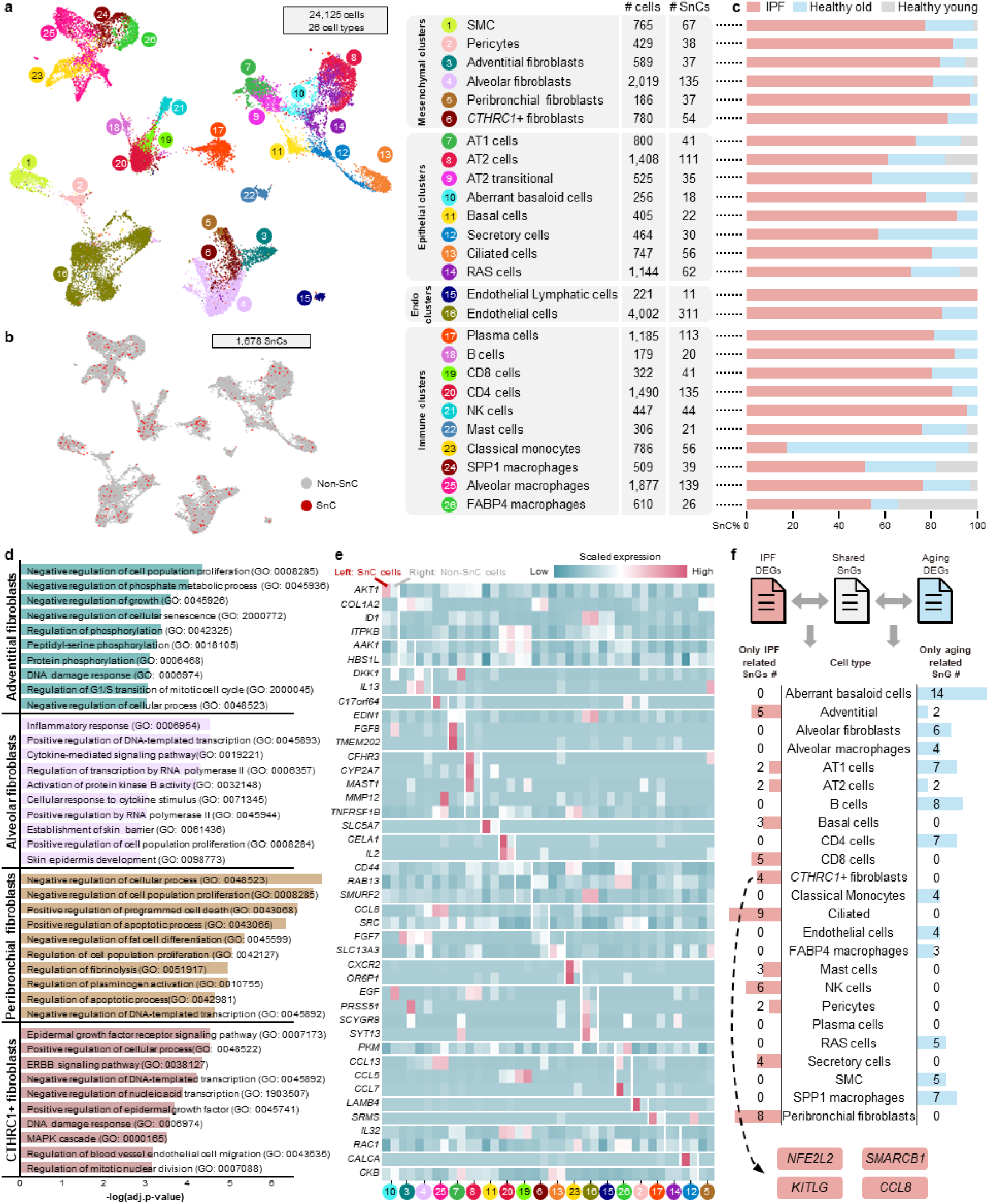
DeepSAS identifies cell-type-specific SnCs and SnGs in IPF lungs. **(a)** UMAP visualization of 24,125 cells from human IPF and healthy lungs, annotated into 25 cell types. The table summarizes the number and proportion of predicted SnCs per cell type identified by DeepSAS. **(b)** UMAP highlighting all DeepSAS-predicted SnCs across the dataset, showing their distribution across multiple cell types. **(c)** SnC percentage of IPF, healthy old, and healthy young samples in each cell type. **(d)** Bar plots showing the top ten enriched GO terms of SnGs in *CTHRC1*+ fibroblast, Peribronchial fibroblasts, alveolar fibroblasts, and adventitial fibroblasts. **(e)** Heatmap of cell-type-specific SnG expression in each cell type. For each cell type, the average expression of SnG in SnCs (left) and non-SnCs (right) is shown. **(f)** Bar plot shows the number of SnGs that are either only related to the IPF phenotype or the aging phenotype. *NFE2L2*, *SMARCB1*, *KITLG*, and *CCL8 were* identified as IPF-related SnGs in *CTHRC1*+ fibroblast.

In parallel, DeepSAS inferred a total of 263 SnGs across the 26 cell types (**Fig. 3d** and **Supplementary Table S7**), highlighting the extensive heterogeneity of senescence programs embedded within different cellular contexts in IPF lungs. Among these, 239 SnGs overlap with at least one of eight curated senescence hallmark gene sets^24–31^, whereas 24 SnGs have not been reported before (**Supplementary Table S8**). Pathway enrichment of SnGs across the four different types of fibroblasts revealed that senescence programs in IPF are strongly shaped by fibroblast identity and niche context rather than representing a uniform cellular state (**Fig. 3d**). Adventitial fibroblasts exhibited enrichment for cell-cycle arrest, phosphorylation signaling, and DNA damage response, consistent with a stress-integrating senescence program in vascular-associated niches. Alveolar fibroblasts showed dominant inflammatory and cytokine-responsive transcriptional activation, reflecting their role in immune crosstalk and injury response within the alveolar compartment. In contrast, peribronchial fibroblasts were enriched for apoptotic regulation and fibrinolysis-related pathways, suggesting a senescence state coupled to epithelial injury and extracellular matrix turnover. Notably, *CTHRC1*+ fibroblasts exhibited enrichment of EGFR, ERBB, MAPK cascade, and DNA damage response pathways, indicating a pathological senescence program that remains transcriptionally active and is closely linked to fibrotic niche remodeling. Such results demonstrate that senescence in IPF manifests as distinct programs aligned with the functional roles and anatomical localization of fibroblast subpopulations, supporting the existence of multiple fibroblast-specific senotypes within fibrotic lungs.

To further examine cell-type specificity, we focused on SnGs uniquely identified within individual cell types. Among the SnGs, 43 were unique to a single cell type (**Supplementary Table S9**), whereas the remaining were shared across two or more cell types. For each cell type, we compared the average expression of its unique SnGs between SnCs and non-SnCs, confirming that these genes exhibited strong and consistent upregulation specifically in senescent cells of the corresponding lineage (**Fig. 3e**). These results demonstrate that DeepSAS captures cell-type-specific transcriptional programs that distinguish SnCs from non-SnCs within the same lineage, rather than reflecting global stress responses shared across cell types.

In addition, the presence of IPF, healthy aged, and healthy young samples in this dataset provides an opportunity to further examine whether the SnGs identified by DeepSAS preferentially associate with disease or aging. To this end, we performed post-hoc analyses^32^ on the 263 DeepSAS-predicted SnGs. Within each cell type, we conducted DEG analysis between IPF and healthy samples and between aged and young samples and intersected the resulting DEGs with the corresponding SnGs. This approach enabled stratification of SnGs into IPF-related, aging-related, shared between IPF and aging, or phenotype-independent categories (**Fig. 3f** and **Supplementary Table S10**). Across cell types, the number and composition of phenotype-associated SnGs varied substantially, underscoring the context-dependent nature of senescence programs. Notably, *CTHRC1*+ fibroblasts exhibited a distinct enrichment of SnGs that were exclusively associated with IPF but not aging. Among these, four genes (*i.e.*, *NFE2L2*, *SMARCB1*, *KITLG*, and *CCL8*) emerged as uniquely IPF-related SnGs in this cell type. *NFE2L2* is a central regulator of oxidative stress responses and redox homeostasis, and increasing evidence indicates that *NFE2L2* can mediate senescence in fibroblasts by inducing a distinct SASP enriched for extracellular matrix and matrisome-associated components^33^, with important roles in wound healing and epithelial cell responses. The enrichment of *NFE2L2* therefore suggests a stress-adaptive, fibroblast-specific senescence state that is closely aligned with the biological function of *CTHRC1*+ fibroblasts in fibrotic niche remodeling. *SMARCB1*, a core component of the SWI/SNF chromatin remodeling complex, implicates transcriptional and epigenetic stabilization of senescence programs that may lock *CTHRC1*+ fibroblasts into a persistent pathological state. In parallel, *KITLG* and *CCL8* reflect non-cell-autonomous outputs of senescence, highlighting the capacity of senescent *CTHRC1*+ fibroblasts to actively modulate their microenvironment through paracrine support and inflammatory cell recruitment.

### Spatial Validation Confirms Cell-Type-Specific SnG Predictions in Fibrotic Lungs

To spatially validate the SnGs predicted by DeepSAS, we analyzed in-house high-resolution spatial transcriptomics data generated using the Xenium platform from one normal lung parenchyma sample and paired upper and lower lobe samples from two IPF patients (**Supplementary Fig. S8**). Altogether, 480 genes were identified in Xenium using our customized gene panel (**Supplementary Table S11**). For IPF samples, lung tissue sections were pathologically annotated into two histologically distinct regions: fibrotic foci and surrounding fibroblast-rich regions. Previous studies have established that alveolar fibroblasts residing in surrounding regions can differentiate into pro-fibrotic *CTHRC1*+ fibroblasts that localize within the fibrotic foci in IPF^34^. Based on this, we used surrounding regions and fibrotic foci as spatial surrogates for alveolar fibroblasts and *CTHRC1*+ fibroblasts, respectively. To quantitatively assess spatial enrichment, we manually annotated 126 fibrotic foci regions (61 upper lobe and 65 lower lobe regions) and 101 surrounding regions (50 upper lobe and 51 lower lobe regions) in IPF. Additionally, we randomly selected 19 regions within the normal parenchyma of the same size to serve as baselines.

We selected *NFE2L2* for validation as a predicted senescence-associated gene in *CTHRC1*+ fibroblasts based on its inclusion in the Xenium panel design. We examined the spatial distribution of *NFE2L2* together with canonical senescence markers and fibroblast lineage markers (**Fig. 4a**). Representative whole-section Xenium images revealed distinct fibrotic architecture, within which *NFE2L2* signal co-localized with *CTHRC1+* fibroblast markers, including *CTHRC1*, *POSTN*, *TGFB1*, and *COL10A1* (**Fig. 4b**, an example of the IPF lower lobe sample). High-magnification views further highlighted enrichment of *NFE2L2* and *CDKN1A* within fibrotic foci compared with surrounding regions. The result showed that *NFE2L2*+ cells were significantly enriched within fibrotic foci compared with surrounding regions and normal parenchyma in both upper and lower lobes (**Fig. 4c**). In contrast, although *CDKN2A* and *CDKN1A* also exhibited higher proportions in fibrotic regions relative to surrounding tissue, these differences reached statistical significance only in the lower lobe and were not consistently observed across lobes (**Fig. 4d-e**). Moreover, *CDKN2A*+ cells were substantially less abundant across all samples compared with *NFE2L2*+ and *CDKN1A*+ cells, indicating the limited sensitivity of this canonical marker in the spatially resolved analyses.

**Fig. 4.**
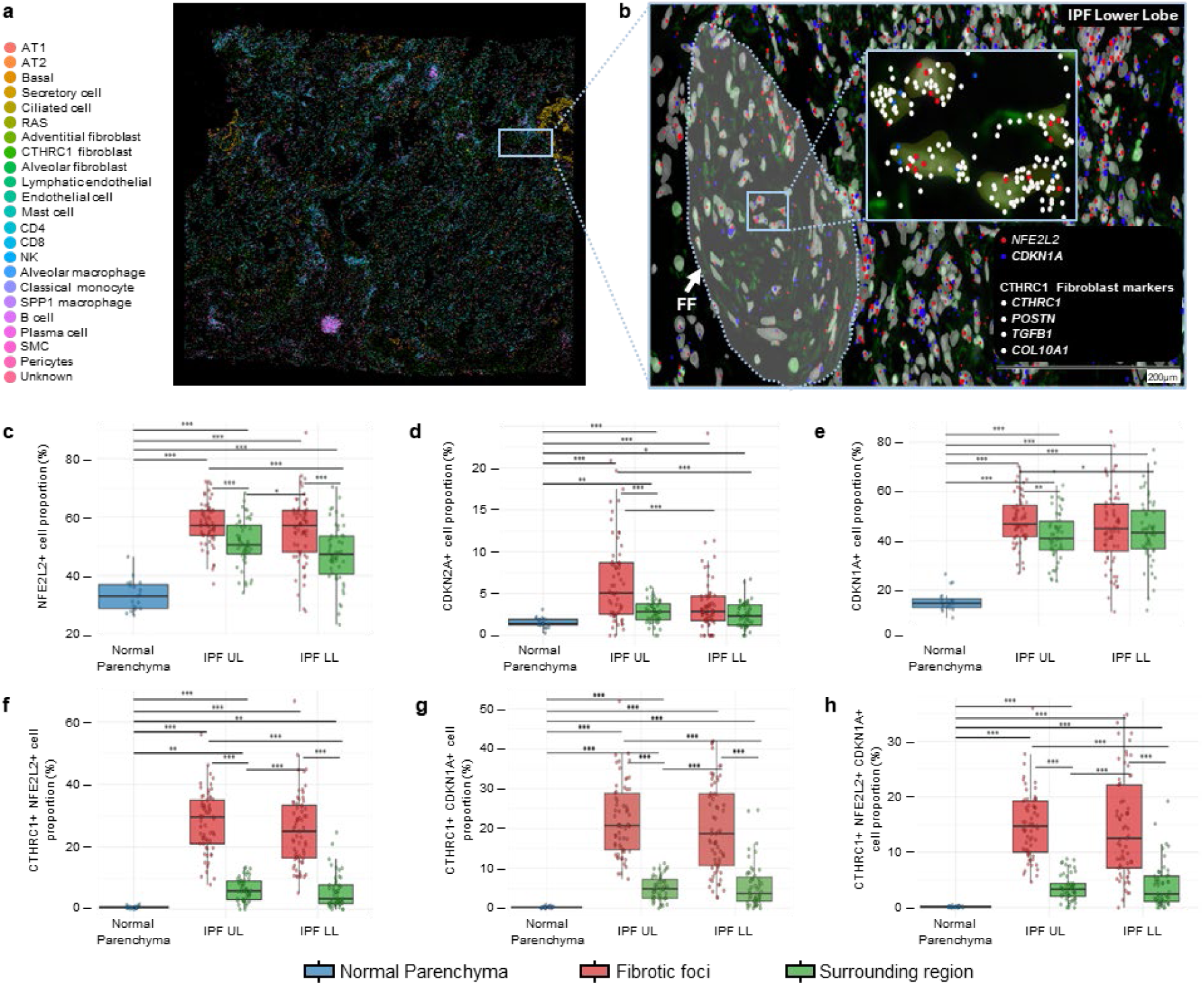
Spatial validation of CTHRC1 fibroblast-specific SnGs in IPF lungs. **(a)** An example of a whole-section Xenium spatial transcriptomics image of the IPF lower lobe lung sample, showing expression of *NFE2L2*, *CDKN1A*, and *CTHRC1*+ fibroblast marker genes, including *CTHRC1*, *POSTN*, *TGFB1*, and *COL10A1*. **(b)** Magnified view of representative fibrotic foci and surrounding fibroblast-rich regions, highlighting spatial enrichment of *NFE2L2* and senescence markers within fibrotic lesions. **(c–e)** Quantification of *NFE2L2*+, *CDKN2A*+, and *CDKN1A*+ cell proportions across selected regions in normal parenchyma, IPF upper lobe, and IPF lower lobe samples. Box plots summarize results from 19 normal regions, 126 fibrotic foci regions, and 101 surrounding regions. Statistical significance was assessed using the Kruskal–Wallis test. *p* < 0.05 (*), *p* < 0.01 (**), and *p* < 0.001 (***) **(f)** Proportion of cells co-expressing *CTHRC1* and *NFE2L2* across spatial regions. **(g)** Proportion of cells co-expressing *CTHRC1* and *CDKN1A* across spatial regions. **(h)** Proportion of triple-positive cells co-expressing *CTHRC1*, *NFE2L2*, and *CDKN1A* across spatial regions.

We next assessed whether restricting the analysis to the pathological fibroblast compartment enhances the specificity of senescence-associated signals. When focusing exclusively on *CTHRC1*+ cells, co-expression of *CTHRC1* and *NFE2L2* was markedly enriched in fibrotic foci compared with surrounding regions in both upper and lower lobes, with substantially greater statistical significance relative to analyses without cell-type restriction (**Fig. 4f**). A comparable, though quantitatively weaker, pattern was observed for the co-expression of *CTHRC1* and *CDKN1A* (**Fig. 4g**).

Furthermore, we examined triple-positive cells co-expressing *CTHRC1*, *NFE2L2*, and *CDKN1A* as a stringent indicator of senescent *CTHRC1*+ fibroblasts. These triple-positive cells were significantly enriched within fibrotic foci relative to surrounding regions in both lobes (**Fig. 4h**), further supporting the preferential spatial localization of senescent *CTHRC1*+ fibroblasts within fibrotic lesions. A similarly weak enrichment was observed for cells co-expressing *CTHRC1*, *NFE2L2*, and *CDKN2A* (**Supplementary Fig. S9**). With these analyses, we confirm that *NFE2L2* represents a *CTHRC1*+ fibroblast-specific, IPF-related SnG, consistent with predictions generated by DeepSAS from single-cell data. The preferential enrichment of *NFE2L2* within fibrotic foci, together with its strong dependence on *CTHRC1*+ fibroblast identity, supports a model in which chronic tissue injury and fibrotic remodeling engage an *NFE2L2*-associated, stress-adaptive senescence state specifically in pathogenic fibroblasts. In contrast to canonical senescence markers (*e.g.*, *CDKN1A* and *CDKN2A*), *NFE2L2* may capture a senescence state that is coupled to a sustained transcriptional and microenvironmental activity within fibrotic lesions, thereby distinguishing disease-associated fibroblast senescence from age-related senescence distributed throughout the tissue. Additional validations were performed for other senescence-associated genes. For example, *CCL4*, identified as a SnG in alveolar fibroblasts, was preferentially enriched in surrounding regions compared with fibrotic foci and was also significantly elevated relative to normal parenchyma. Consistent with a senescent phenotype, cells co-expressing *CCL4* and *IL6*, a canonical marker of senescence, were significantly more abundant in surrounding regions compared with fibrotic foci. (**Supplementary Fig. S10**). Taken together, these findings provide orthogonal validation that DeepSAS identifies biologically meaningful and spatially coherent senescence programs embedded within pathological fibroblast populations in IPF lungs.

### Senescence Programs Differ Substantially between IPF lungs and bleomycin-treated hPCLS

Bleomycin treatment is widely used as an exogenous trigger of DNA damage and tissue injury, inducing lung fibrosis in mice and senescence in human ex vivo models such as hPCLS. hPCLS are generally considered to recapitulate key aspects of the native lung architecture and cellular proportions. Accordingly, hPCLS is frequently employed as a translational model to interrogate and validate senescence-associated mechanisms observed in lung diseases^35^. Based on this premise, we sought to assess whether senescence programs identified in IPF lungs could be reproduced in healthy human hPCLS samples treated with bleomycin. As expected, the comparison of untreated control and bleomycin-treated hPCLS samples revealed changes in cell-type composition that resemble the IPF lung. These changes include loss of Alveolar type 2 and type 1 epithelial cells, reduced pericytes and NK cells, with a parallel increase in aberrant basaloid epithelial cells and inflammatory and *CTHRC1*+ fibroblasts (**Supplementary Fig. S11**).

We applied DeepSAS to the hPCLS model that included both untreated control and bleomycin-treated samples (**Fig. 5a**). In general, DeepSAS revealed markedly stronger senescence signals in acutely injured bleomycin-treated hPCLS samples compared with chronic injured IPF lungs. For instance, the proportion of SnCs within the pathological fibroblast *CTHRC1+* population was substantially higher in bleomycin-treated hPCLS than in IPF patient samples, indicating that acute injury elicits a more pronounced senescence response than that observed in chronic disease (**Fig. 5b**). Consistent with this observation, the composition of SnGs differed strikingly between the two contexts. In *CTHRC1*+ fibroblasts, DeepSAS identified 79 SnGs exclusively in bleomycin-treated hPCLS samples, and 36 SnGs exclusively in IPF lungs, with only 19 SnGs shared between the two conditions (**Fig. 5c** and **Supplementary Table S12**). This limited overlap demonstrates that senescence programs triggered by acute bleomycin exposure are largely distinct from those operating in chronic IPF, despite superficial similarities in fibrotic pathology. To further characterize these differences, we performed pathway enrichment analysis separately for the 98 *CTHRC1*+ fibroblast SnGs identified in hPCLS and the 55 SnGs identified in IPF (**Fig. 5d** and **Supplementary Table S13**). Although both conditions showed enrichment for canonical senescence-related processes such as DNA damage response, oxidative stress, mitochondrial dysfunction, inflammatory signaling, and apoptosis, these pathways were consistently more significant in hPCLS. In particular, oxidative stress response, DNA damage signaling, and metabolic reprogramming pathways exhibited markedly stronger enrichment in hPCLS than in IPF, suggesting a more intense and coordinated senescence program in the acute injury setting. In contrast, IPF-associated SnGs showed comparatively attenuated enrichment, consistent with a chronic, lower-intensity senescence state.

**Fig. 5.**
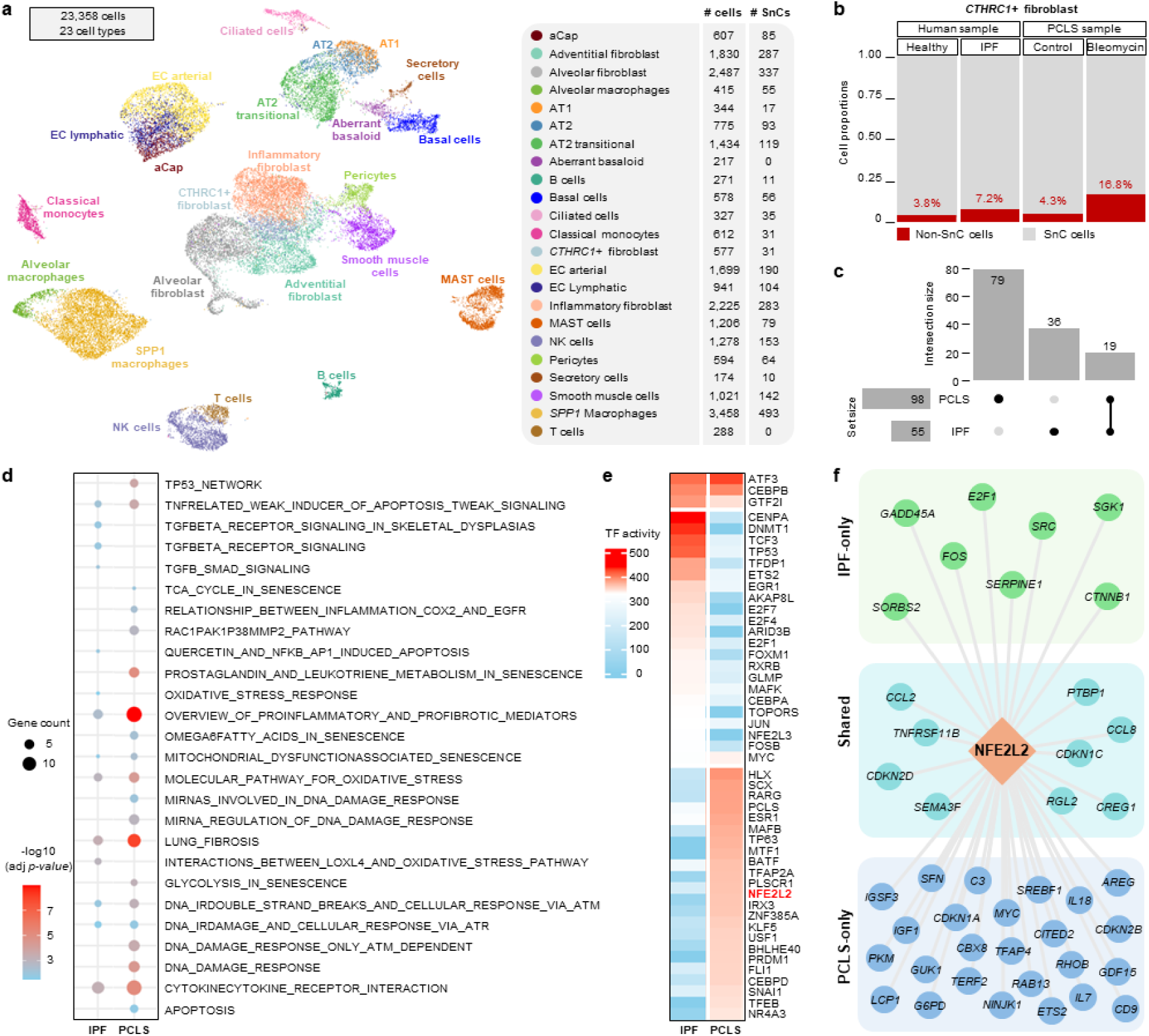
Acute and chronic senescence programs exhibit distinct regulatory features in lung fibroblasts. **(a)** UMAP of the hPCLS scRNA-seq dataset comprising 23,358 cells annotated into 23 cell types. **(b)** Proportion of SnCs in *CTHRC1*+ fibroblast in hPCLS and IPF datasets. **(c)** DeepSAS identified SnGs in *CTHRC1*+ fibroblast in hPCLS and IPF datasets, respectively. **(d)** Pathway enrichment analysis of SnGs identified in hPCLS and IPF, highlighting shared and context-specific senescence-associated pathways. **(e)** Top 15 TF inferred from *CTHRC1*+ fibroblast SnGs in hPCLS and IPF datasets, respectively. TFs are ranked by TF activity scores from high to low and grouped into shared, IPF high, and hPCLS high groups. *NFE2L2* was highlighted in red. **(f)** A *NFE2L2* gene regulation network showing the distinguished and shared genes regulated by *NFE2L2* in IPF and hPCLS data.

We next examined transcription factor (TF) regulation underlying these context-specific senescence programs. TF activity analysis revealed both shared and distinct regulators between bleomycin-treated hPCLS samples and IPF lungs (**Fig. 5e** and **Supplementary Table S14**). Among these, *NFE2L2* emerged as a prominent regulator with substantially higher inferred activity in hPCLS than in IPF, whereas several additional regulators involved in stress response and cell-cycle control also differed between the two contexts. To further dissect the role of *NFE2L2*, we quantified the number of SnGs inferred to be regulated by *NFE2L2* in each condition (**Fig. 5f**). *NFE2L2* regulated a larger set of SnGs in hPCLS than in IPF, including genes involved in oxidative stress response, extracellular matrix remodeling, and secretory signaling. In contrast, the smaller *NFE2L2*-regulated SnG set in IPF suggests a reduced or constrained role of this regulator in chronic injury.

The above findings highlight fundamental differences between bleomycin-induced and IPF-associated senescence. Bleomycin is a potent inducer of oxidative stress and robustly activates *NFE2L2* as an acute protective response, consistent with prior studies showing exacerbated inflammation and fibrosis in *NFE2L2*-deficient bleomycin models^36^. Paradoxically, sustained or hyperactivated *NFE2L2* signaling, particularly in fibroblasts, has been shown to promote fibroblast senescence, SASP enriched in ECM components, and gene set enrichment for pathways promoting epithelial cell activation, cell movement, cytoskeletal remodeling, angiogenesis, and fibrosis. Accordingly, the amplified *NFE2L2*-driven senescence program observed in bleomycin-treated hPCLS represents an acute, stress-induced senescence state that differs fundamentally from the more restrained, chronic senescence observed in IPF patient lungs. Together, these results demonstrate that although bleomycin-treated hPCLS captures key features of senescence, it does not faithfully recapitulate the regulatory architecture of chronic IPF-associated senescence. The limited overlap of SnGs between hPCLS and IPF and the divergent role of *NFE2L2* underscore fundamental differences between acute injury–induced and chronic disease– associated senescence. DeepSAS sensitively resolves these context-dependent senescence programs, highlighting the importance of distinguishing acute and chronic senescence mechanisms when interpreting experimental models of fibrotic disease.

## DISCUSSION

DeepSAS is an AI-driven framework to resolve cellular senescence at single-cell resolution while preserving cell-type specificity. Its core innovation lies in modeling senescence as a cell-type-embedded and context-dependent state rather than a global phenotype. By constructing a heterogeneous cell–gene graph that integrates intracellular transcriptional states, intercellular CCC signals, and external senescence knowledge, DeepSAS enables joint inference of SnCs and SnGs without imposing a predefined order of identification. Attention-based graph learning provides a robust foundation for senescence scoring that is less sensitive to transcriptional sparsity than expression-based activity scores, while an iterative contrastive learning scheme operating within individual cell types allows senescence programs to emerge in a self-supervised manner. Importantly, senescence hallmarks are treated as guidance rather than hard restrictions, enabling the discovery of novel, cell-type-specific SnGs beyond canonical markers.

Across multiple biological contexts, DeepSAS consistently identified biologically coherent senescence programs. In healthy human eye and lung datasets, SnC abundance increased with aging in a cell-type-specific manner, establishing aging as an independent validation axis. In IPF lungs, DeepSAS resolved extensive senescence heterogeneity across cellular compartments and uncovered disease-associated SnGs without incorporating phenotype information during model inference. Spatial transcriptomics further validated these predictions, demonstrating that *NFE2L2* is specifically enriched in senescent *CTHRC1+* fibroblasts within fibrotic foci, with greater cell-type specificity and spatial consistency than canonical markers such as *CDKN1A* and *CDKN2A*. Comparison of a chronic condition (*i.e.*, IPF) with an acute injury model (bleomycin-treated hPCLS) revealed that senescence programs differ substantially between these contexts. In hPCLS, bleomycin-induced senescence showed a distinct and more robust senescence signature, a different SnG composition, and enhanced *NFE2L2* regulatory activity. These findings indicate that acute oxidative injury triggers a senescence state that is not equivalent to the chronic, IPF-associated form of senescence and is instead driven by unique regulatory programs governing senescence-related genes.

Nevertheless, several limitations should be noted. First, DeepSAS currently models senescence as a binary state and does not explicitly capture intermediate or progressive senescence stages. Second, phenotype associations for SnGs are inferred post hoc through DEG analysis rather than intrinsically modeled within the learning framework. Third, although DeepSAS performs robustly across large and diverse datasets, its sensitivity to extreme data sparsity and low-quality clinical samples warrants further benchmarking. Finally, while CCC information improves cell-type specificity, additional sources of spatial or microenvironmental information may further enhance resolution.

Future extensions of DeepSAS will focus on expanding senescence characterization beyond transcriptomics. Integration of multi-omics data, particularly metabolomic and mitochondrial readouts, may provide earlier and more mechanistic indicators of senescence states. Systematic benchmarking across experimental models and disease contexts will further clarify how acute and chronic senescence differ at regulatory and functional levels. More broadly, the ability of DeepSAS to resolve cell-type-specific and context-dependent senescence programs supports a shift toward senotypes as the fundamental unit of senescence biology. Defining senotypes across tissues and diseases will be critical for the development of precision senotherapy^5,6^, enabling targeted intervention strategies that account for both cell type and senescence state rather than relying on universal senescence markers.

## METHODS

### 1. The DeepSAS framework

#### 1.1. Data preprocessing

A raw expression matrix was preprocessed through necessary quality control processes, including cell filtering, gene filtering, doublets detection and removal, batch removal, normalization, and scaling. The processed data were integrated for joint clustering and cell type annotation. High-quality clustering and accurate annotation results can enhance the SnC and SnG prediction in DeepSAS. In the absence of cell type labels, DeepSAS employed the standard Louvain clustering algorithm^37^ to categorize cells into distinct clusters.

#### 1.2. Heterogeneous graph construction

The preprocessed expression matrix is defined as X^𝑅^ ∈ ℝ^𝑁×𝑀^, where 𝑁 denotes the number of cells and 𝑀 denotes the number of genes. We constructed an undirected cell-gene heterogeneous graph 𝐺 = (𝑉, 𝐸) from X^𝑅^ , incorporating the node set 𝑉 = 𝑉^𝐶^ ∪ 𝑉^𝐺^ where 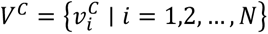 represent cells, and 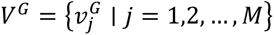 represent genes. By default, DeepSAS utilizes all genes in the following steps to ensure that potentially lowly expressed SnGs can be captured. To improve computational efficiency, we also offer the option to select only the top highly variable genes. The edge set 𝐸 comprises cell-gene edge 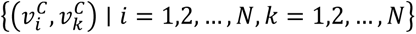 and cell-cell edge 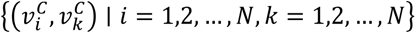 . The cell-gene edge 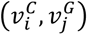 represents the gene expression of 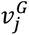 in 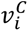. For 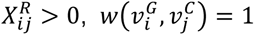, otherwise, 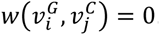. The cell-cell edge 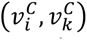 represents the CCC between 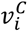 and 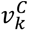, in terms of SASP-related ligand-receptor interaction probability predicted by SoptSC^38^. Differing from other existing CCC prediction tools, SoptSC calculates the probability of CCC between two cells, rather than two cell types. Suppose we have a ligand-receptor pair 𝑝_𝑘_ , with expression distribution given by two vectors 𝐿^𝑘^ ∈ ℝ^𝑁^ and 𝑅^𝑘^ ∈ ℝ^𝑁^ for ligand and receptor, respectively, the probability that a signal is sent from cell 𝑖 to cell 𝑗𝑗 by this ligand-receptor pair 𝑝_𝑘_ is then given by:

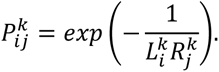

We then constructed the CCC probability matrix 𝑃^𝑘^ ∈ ℝ^𝑁×𝑁^ regarding the ligand-receptor pair 𝑝_𝑘_. The overall probability of signaling summed over all ligand-receptor pairs, 𝑃_1_ ∈ 𝑅^𝑁×𝑁^, is given by

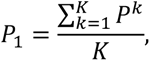

where 𝐾 is the number of all ligand-receptor pairs. 𝑃_1_ serves as the adjacency matrix of the CCC graph. We symmetrized this matrix by considering bidirectional interactions:

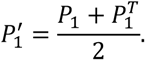

Thus, the edge weight between two cells in the heterogeneous graph is defined based on a threshold 𝜙 for signaling strength:

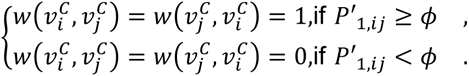

Lastly, we computed initial embeddings for genes and cells via UMAP dimensionality reduction. The UMAP algorithm processes the cell-gene expression matrix X^𝑅^, generating a low-dimensional embedding matrix 𝑍_𝑐_ ∈ ℝ^𝑁×𝐷^ for each cell, where 𝐷 is the dimension of the embedding vector. The expression matrix is then transposed, and UMAP is applied to generate a low-dimensional embedding matrix for genes 𝑍_𝑔_ ∈ ℝ^𝑀×𝐷^. Thus, each cell and gene obtains the same initial embedding of 𝐷 dimensions. The final embedding matrix 𝑍^ ∈ ℝ^(𝑀+𝑁)×𝐷^ represents the embedding for all genes and cells.

#### 1.3. Graph autoencoder

We adopted a graph autoencoder to learn graph embeddings for all the nodes and edges in 𝐺 defined in the above section. The graph autoencoder aims to reconstruct the adjacency matrix 𝐴 of the heterogeneous graph 𝐺. It consists of an encoder layer and a decoder layer, similar to a standard autoencoder. The encoder layer of the graph autoencoder utilizes a graph attention network (GAT), while the decoder layer is based on the inner product of the latent embeddings. The inputs to the encoder layer are the embedding matrices 𝑍_𝑐_ for cell nodes and 𝑍_𝑔𝑔_ for gene nodes, along with the adjacency matrix 𝐴. Due to the heterogeneity of nodes, different types of nodes reside in distinct feature spaces. Therefore, for each type of node (e.g., node with type 𝜙_𝑖_), we designed a type-specific transformation matrix 𝑀_𝜙𝑖_ to project the features of different node types into a common feature space:

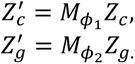

The encoder layer comprises 𝐿 GAT layers. The embedding of node 𝑖 in the 𝑙-th layer is 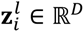, and the updated embedding of node 𝑖 in the (𝑙 + 1)-th layer is:

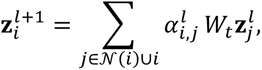

where the attention coefficients 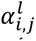 are computed as

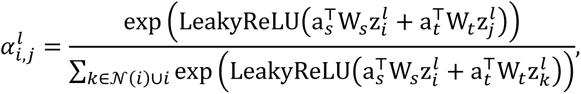

where W_𝑠_ and W_𝑡_ are the weight matrices for the source node and target node, respectively. When a cell is treated as a target node, its connected genes and neighboring cells serve as source nodes, similarly to how a gene acts when selected as the target. A is the weight vector of attention, 𝒩(𝑖) is the set of neighbor nodes of the node 𝑖, 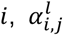 is the attention coefficient between node 𝑖 and 𝑗. The outputs of the encoder layer are the embedding matrix of cells Z_𝑐_ ∈ ℝ^𝑁×𝐷^^1^ and genes Z_𝑔𝑔_ ∈ ℝ^𝑀×𝐷^^1^ . Then the reconstructed adjacency matrix *Â*^′^ is calculated by:

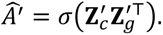

𝜎 is a sigmoid function. The loss function of the graph autoencoder is defined as the binary cross-entropy between the estimated and the ground-truth edges:

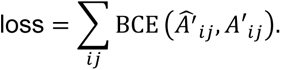

#### 1.4. Reconstruct the heterogeneous graph

To support biologically guided contrastive learning, we reconstruct a weighted heterogeneous bipartite graph 𝐺^′^ = (𝑉, 𝐸^′^), in which edges represent connections derived from the graph transformer in Section 1.3. We define the edge set 𝐸^′^ ⊆ 𝑉^𝐶^ × 𝑉^𝐺^, where each edge 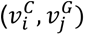 is assigned a weight 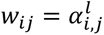with 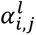 the attention score from the final heterogeneous graph transformer layer 𝑙. Each gene node is assigned to a binary label as follows:

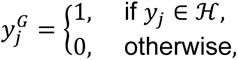

where ℋ is the union of senescence hallmark genes (e.g., SenMayo^28^, Fridman^29^, CellAge^30^) and cell-cycle markers. Each cell node is assigned a discrete label 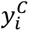 based on the cell type or cell cluster labels defined in 1.1 Data preprocessing. Lastly, we retain the encoded latent embeddings of all nodes from the encoder output of the graph autoencoder.

#### 1.5. Contrastive learning iteration

This framework iteratively identifies candidate SnGs and SnCs. During the iteration process, we let the encoder 𝑓_𝜃_ learn how to distinguish between the SnCs and non-SnCs by maximizing the distance between them. Initially, the known senescent marker lists determine the senescent gene nodes. We defined a senescence score attention score 𝛼_𝑖_ of cell 𝑖 based on the average attention scores of SnG candidates:

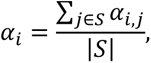

where 𝑆 is the set of SnG candidates, |𝑆| is the total number of SnG candidates. We select the top-𝐷_1_ cells with the highest senescence score as SnCs in the first iteration. 𝐷_1_ is defined by the upper bound of the senescence score distribution, 𝑄_3_ + 1.5 × IQR, where 𝑄_3_ is the third quartile, and IQR is the interquartile range. We retain cell types with more than 10 senescent cells.

We next calculated senescence scores for genes. Similar to the selection of SnC candidates, we computed, for each gene, the average attention scores across SnC candidates within the same cell type. The top-𝐷_2_ genes with the highest senescence scores were selected, where 𝐷_2_ is defined as the upper bound of the senescence score distribution for SnG candidates. That is to say, starting from the second iteration, each cell type maintains its own SnG candidate list based on these senescence scores.

We defined three different distances 𝑑_1_, 𝑑_2_, 𝑑_3_ with biological significance to calculate the distance between the cell nodes. 𝑑_1_ is the Euclidean distance between a SnC and the center of the SnC hub in the same cell type. 𝑑_2_ is the distance between two SnCs in different cell types. 𝑑_3_ is the distance between a SnC and a non-SnC in the same cell type. We hypothesize that the relationship should satisfy the following conditions:

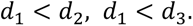

To enforce these constraints, the loss function ℒ is defined as follows:

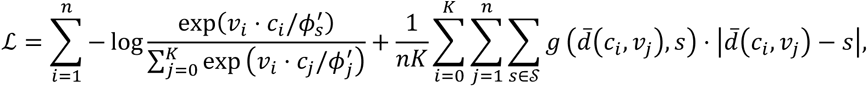

where 𝑣_𝑖_ is the embedding for cell node 𝑖, 𝑐_𝑖_ is the embedding of the center of the cluster where node 𝑖 belongs to. 𝐾 is the number of cell clusters. 𝑛 is the total number of cells. The first term is the ProtoNCE loss function^39^. ProtoNCE could estimate the concentration for the feature distribution around each cell cluster. The distribution of embeddings for each cell cluster has a different level of concentration. We use 𝜙^′^ to denote the concentration level of the feature distribution:

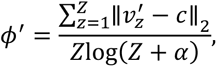

where 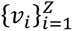 are the momentum features and 𝑐 is the hub center of SnCs in one cell type. 𝛼 is a smooth parameter to ensure that small clusters do not have an overly large 𝜙^′^. A smaller value of 𝜙^′^ indicates a larger concentration. The desired 𝜙^′^ value should be small (i.e., high concentration) if (*i*) the average distance between 𝑣^′^ and 𝑐 is small, and (*ii*) the cluster contains more feature points (i.e., 𝑍 is large). 𝜙^′^ is then normalized for all clusters. The second term in the loss function is the Multi-level Distance Regularization (MDR)^40^. The distance is defined as the normalized Euclidean distance between two given embedding vectors:

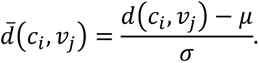

We define a set of levels 𝑠 ∈ 𝖲, and the levels are initialized with predefined values; each level 𝑠 is interpreted as a multiplier of the standard deviation of the normalized distance. 𝑔(𝑑; 𝑠) is an assignment function that outputs whether the given distance *d̂* and the given level 𝑠 are the closest or not, and is defined as:

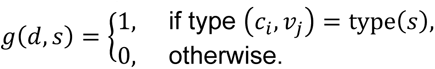

By adopting the assignment function, MDR selects valid regularization levels for each distance with consideration of various degrees of similarity. After optimizing with the loss function ℒ, the cell embeddings are updated. The iteration converges when the overlap of candidate SnCs and SnGs between adjacent iterations reaches 1. The output of this step is a SnC label (0 represents non-SnC and 1 represents SnC) for each cell, an attention-based senescence score for each cell, and an unfiltered SnG list in each cell type.

#### 1.6. Cell type shared and specific SnG determination

To determine SnGs, we applied a three-level selection strategy.

- The first level consists of the direct output of the contrastive learning framework. For predicted SnCs in each cell type, the adjacent genes with the highest senescence scores were selected.
- To enhance experimental interpretability, we next assessed expression differences within each cell type by grouping cells into SnCs and non-SnCs and performing differential expression analysis. SnG in each cell type identified in the last step was then intersected with the corresponding DEGs (log fold change ≥ 0.25) within the same cell type to ensure transcriptional distinction.
- Finally, to capture cell-type-specific signals, we retained only those genes from the second set that appeared exclusively in a single cell type. These were designated as cell-type-specific SnGs. Genes that appeared in more than one cell type after DEG filtering were referred to as cell-type-shared SnGs.

This hierarchical strategy ensured that final SnGs are not only biologically relevant and discriminative between SnCs and non-SnCs, but also practical candidates for downstream validation within specific cellular contexts. Using the same strategy, we also evaluated the relationship between SnGs and sample phenotypes. Specifically, within each cell type, we stratified cells by either age group (young vs. old) or disease status (healthy vs. IPF) and performed differential expression analysis. The resulting DEGs were then intersected with the cell-type-shared SnGs to identify age-related SnGs and IPF-related SnGs, respectively. This allowed us to further characterize the phenotypic relevance of senescence programs across distinct cellular contexts.

### 2. Evaluation and benchmarking

#### 2.1. Robustness test

To assess the robustness of DeepSAS against variations in dataset composition, we conducted a series of subsampling experiments designed to simulate real-world scenarios with limited or imbalanced data. The full dataset (comprising IPF and healthy control samples) served as the reference for comparison. We generated multiple modified versions of the data by applying strategic subsampling or removal procedures and evaluated how consistently DeepSAS could recover SnCs and associated predictions. Specifically, we performed ten distinct subsampling schemes. For each, we randomly selected a fixed fraction (2/3, 1/2, and 1/3) of cells from the healthy samples and IPF samples, either jointly or separately. In total, six subsampling conditions were created:

- two-thirds of the cells from both healthy and IPF samples
- one-half from both
- one-third from both
- two-thirds from only IPF samples
- one-half from only IPF samples
- one-third from only IPF samples

In addition, to evaluate robustness to missing cellular contexts, we performed a cell-type dropout test: in each run, we randomly selected 10 out of the 25 annotated cell types and removed all cells from those types.

Each of the above subsampling scenarios was repeated five times to ensure consistency. DeepSAS was independently applied to each replicate. The set of SnCs identified in each subsampled run was compared to the original SnCs predicted from the full dataset. We recorded the overlap rate and number of overlapping SnCs between the subsampled and reference results to quantify stability.

#### 2.2. Ablation test

To dissect the contribution of individual components in the DeepSAS framework, we conducted an ablation study by systematically modifying or removing specific modules. Each variant was benchmarked against the full DeepSAS model with default settings to evaluate its impact on SnC identification performance. First, we examined the role of CCC by disabling the CCC-based edge augmentation in the graph construction step. This allowed us to assess the importance of incorporating predicted intercellular signaling into the heterogeneous graph structure. Second, we tested the effect of node set size by varying the number of genes included per cell. Specifically, we reduced the default gene set to subsets of 8,000 and 3,000 genes, simulating conditions with limited gene coverage or lower sequencing depth. Third, we evaluated the sensitivity of DeepSAS to different senescence hallmark gene sets used to initialize senescence-associated gene (SnG) candidates. We tested four alternatives: (i) Fridman only, (ii) SenMayo only, (iii) SenMayo + CellAge, and (iv) SenMayo + Fridman. Each configuration was run independently, and outputs were compared to the default multi-source initialization. Performance metrics, including the number and distribution of identified SnCs and SnGs and overlap with reference results, were used to quantify changes in model behavior. This ablation study highlights the relative contributions of key architectural choices and confirms the robustness and interpretability of DeepSAS under varying input assumptions and configurations.

#### 2.3. Comparison with other tools

To evaluate DeepSAS in the context of senescent cell identification, we performed a benchmarking analysis against several contemporary computational approaches selected for their focus on rare cell populations, transcriptomic heterogeneity, or senescence-specific modeling.

- ***SenCID*** trains a support vector machine using published bulk RNA-seq data from both normal and SnCs^9^. One limitation of SenCID is that its training data were collected from in vitro studies, which cannot be transferred to in vivo conditions. Also, it often predicts an unreasonably high proportion of cells (>70%) as senescent in every cell type, based on our tests, suggesting poor specificity and possible overfitting to general stress responses. Rather than identifying cell-type-specific SnGs, SenCID identifies six senescence gene sets to cover senescence characterization in 30 cell types, yet it is not capable of generating new SnGs for new input scRNA-seq data.
- ***SenePy*** separates the derivation of senescence gene signatures from cell scoring, leading to biased or suboptimal identification of SnCs and SnGs^10^. Its reliance on co-expression and binarized networks makes it sensitive to dropout, likely missing lowly expressed or transient markers. Moreover, our team tested the SenePy tool and found that it cannot generate new cell-type-specific SnGs from a new dataset, lacking the discovery power needed to derive cell-type-specific SnGs directly from input scRNA-seq data.
- ***DeepScence*** lacks cell-type specificity, relies on self-compiled senescence hallmarks for training, and does not infer SnGs^11^. Its binary classification approach may also miss subtle or context-specific senescence phenotypes. These constraints position DeepScence as less flexible and less biologically contextualized. Since this work has not yet been published, we did not include it in the benchmark.
- **Seurat+ChatGPT**. We adapted a strategy inspired by the GPTCelltype framework^41^, using Seurat for differential expression analysis alongside large language model (LLM)-assisted annotation. Specifically, for each cluster identified within the dataset, we extracted the top 20 DEGs using Seurat. These DEGs were embedded into a structured prompt—“This is a list that includes senescent cell marker:[*Genelist*] I have [*number*] clusters, some of them are senescent cells, provide the accurate cell names with whether they are senescent cell and the senescent biomarkers you use to do senescent dentification the gene markers are found here: (1) Differential expressed genes for group 0: [*Genelist*]; (2) Differential expressed genes for group 1: [*Genelist*]…”—which we input to ChatGPT-4. The resulting annotations leveraged both the statistical power of Seurat’s clustering and the contextual reasoning provided by LLM to discern potential senescent subsets.
- **MarsGT+ChatGPT**. Because senescent cells often represent a minority population, we additionally tested MarsGT^42^, a method specifically tailored to detect rare cell types. MarsGT was first used to cluster the transcriptomic dataset, after which we applied Scanpy to identify the top 20 DEGs in each rare cell cluster. These gene lists were then incorporated into the same GPT prompt template described above and analyzed via ChatGPT-4 to further refine senescence-related cluster assignments.

### 3. Case study 1: healthy human retinal pigment epithelium and choroid snRNA-seq data

snRNA-seq data from the human RPE and choroid were obtained from the posterior segments of postmortem eyes of 53 donors in Dr. Rui Chen’s lab. Nuclei were isolated, and snRNA-seq libraries were generated using the Chromium Next GEM Single Cell 3′ Reagent Kit v3.1 (10x Genomics) and sequenced on the NovaSeq 6000 platform (Illumina). Data processing followed the methodology previously described in a preprint by Jin Li et al.^43^. Briefly, sequencing reads were aligned using Cell Ranger (v7.1.0, 10x Genomics), and quality control was performed using cellqc (https://github.com/lijinbio/cellqc) with the following criteria: correction for ambient RNA contamination; removal of low-quality nuclei with fewer than 300 detected features, fewer than 500 total transcript counts, or with >10% of reads mapping to mitochondrial genes; and doublet detection using DoubletFinder. Batch correction and data integration across multiple samples were conducted using scVI^44^. Major cell types were annotated based on canonical marker genes. The dataset comprising 716,410 single cells was randomly subsampled, but preserved the same cell proportion into 29 batches. The first 28 batches involve 25,000 cells per batch, and the final (29^th^) batch included the remaining 16,410 cells. Each batch was independently analyzed using DeepSAS.

### 4. Case study 2: healthy human lung scRNA-seq data

scRNAseq data from healthy human fetal lung samples were obtained from the Gene Expression Omnibus under accession GSE260769. Raw count matrices were downloaded and processed using Seurat (v4). For each developmental time point, a Seurat object was created from the 10x Genomics output using *CreateSeuratObject*, and mitochondrial read content was quantified as the percentage of transcripts mapping to mitochondrial genes. Quality control was performed separately for each sample, retaining cells with more than 200 and fewer than 7,000 detected genes and with mitochondrial content below 10%. Following quality filtering, datasets were normalized independently using log normalization, and highly variable genes were identified within each sample. Integration features were selected across all samples, after which the data were scaled, and principal component analysis was performed using the shared feature space. Cell types were annotated manually based on the original annotations provided in the source publication and further refined using canonical marker gene expression. The dataset comprising 45,370 single cells was randomly subsampled, but preserved the same cell proportion into 5 batches. Each batch was independently analyzed using DeepSAS.

### 5. Case study 3: in-house human IPF single-cell Flex and Xenium data

#### 5.1. Sample collection and processing

Collection of human lung tissue was approved by the Institutional Review Board (IRB) of The Ohio State University (OSU) under study ID 2021H0180 and was performed by the Comprehensive Transplant Center (CTC) Human Tissue Biorepository at OSU with Total Transplant Care Protocol informed consent and research authorization from the donors. CTC operates in accordance with NCI and ISBER Best Practices for Repositories. The demographic summary of the study population is shown in **Supplementary Table S15.** Lung tissues were collected from the upper and lower lobes of lung explants of patients diagnosed with IPF and from the parenchymal anatomical region of age-matched and young non-IPF donors. The lung tissue was minced into pieces of 10 mm x 20 mm and fixed for 24 h in 4% paraformaldehyde and then transferred to 70% ethanol and embedded in paraffin (wax) to create a Formalin-Fixed Paraffin-Embedded (FFPE) lung tissue block.

#### 5.2. Flex scRNA-seq data generation and analysis

##### 10X Flex scRNA-seq library preparation

Fixed RNA Profiling was performed on one biological sample and two biological samples from the parenchymal lung region of young non-IPF and age-matched normal donors, respectively, and on two biological samples from the parenchymal lung region of upper and lower lobes from two unique IPF patients. For each donor or explant, two FFPE blocks were obtained and preprocessed to evaluate lung architecture (H&E staining) and RNA quality (parameter >30% DV200) to select one FFPE block per sample. For each sample, two tissue sections at 50 µm were then cut from the FFPE blocks and placed onto 1.5 ml PCR clean tube. Then, from cells isolated from FFPE tissue using 1 mg/ml Liberase TH (Dissociation protocol, CG000632, Rev D, 10x genomics), the fixed RNA profiling and library preparation (Chromium Fixed RNA Profiling Reagent Kits for Multiplexed Samples, CG000527, Rev F, 10x Genomics) were performed based on human probes to identify whole transcriptome, with a target of 128,000 cells for capture, 8,000 cells per sample for a single reaction of 16 samples, each sample with a barcode. Next-generation sequencing was carried out in the Advanced Genomics Core at the University of Michigan and in the Novogene Corporation Inc. Gene expression libraries were sequenced on Illumina NovaSeq 6000 sequencer with sequencing depth of 15,000 read pairs per cell, Paired-end, dual indexing type, read lengths of 28 cycles Read 1, 10 cycles i7 index, 10 cycles i5 index, 90 cycles Read 2. scRNA-seq data was extracted from the raw sequencing data using Cell Ranger (version 7.1.0, 10x Genomics).

##### IPF raw sequence mapping and quantification

Cell Ranger^45^ (version 2.1.0, 10x Genomics) was used for read alignment, transcript reconstruction/annotation, abundance estimation, read quality trimming, and sample quality analysis. Background expression due to ambient RNA was mitigated using CellBender^46^, while doublets were identified using DoubletFinder^47^. The output gene expression file was analyzed using Seurat v4.4.0^48^. Low-quality cells were identified and removed based on high mitochondrial (>10%) and low transcriptional counts (<200). The preprocessed expression matrix was then used for dimensional reduction and cell clustering using Seurat. DEGs between different clusters were determined by the Wilcoxon Rank Sum test, where p-values were adjusted for multiple testing using the Bonferroni procedure. Clusters were annotated by comparing DEGs to canonical markers from the public domain (**Supplementary Table S5**).

#### 5.3. Xenium data generation and validation

##### Sample preparation on Xenium Slides

FFPE-lung tissue was sectioned 5×6 mm at 5 µm onto a Xenium slide. Sections were treated to access the RNA for labeling with circularizable DNA probes.

##### Gene panel design

Xenium in situ technology relies on a pre-defined gene panel. Each probe consists of two complementary sequences targeting the mRNA of interest and a unique gene-specific barcode. Upon hybridization, the probe circularizes and undergoes rolling circle amplification, amplifying the signal for target detection and decoding. A total of 480 genes were derived from a standalone human lung Xenium panel (ID3ZXPF2) designed by the Lung Aging Lab and analyzed using Chemistry version v1 on the Xenium Analyzer instrument serial number XETG00239. The custom panel was curated based on human lung single-cell analysis data generated by the Lung Aging Lab (**Supplementary Table S11**), focusing on genes relevant for cell type identification and potential involvement in aging, cellular senescence, and IPF. Finally, a spatial map of the transcripts in the entire tissue section was built using Xenium analysis software version 2.0.0.10 and instrument software version 2.0.1.0, which allowed immediate exploration of the assay’s subcellular readout.

##### Cell segmentation and annotation

The Xenium In Situ Cell Segmentation Kit (Xenium Multi-Tissue Stain), PN-1000662 (10x genomics) was used to improve the determination of cell boundaries by using a stain and algorithmic technique developed and validated through custom-trained machine learning models. We manually annotated cell clusters and confirmed that Xenium spatial transcriptomics captured the heterogeneous cellular populations within lung tissue. Alveolar type 1 (AT1) cells were identified by the expression of *CLIC5*, *AGER*, and *RTKN2*, while alveolar type 2 (AT2) cells were characterized by *SFTPC*, *LAMP3*, and *HHIP*. Basal cells exhibited high expression of *KRT17*, *TP63*, and *KRT5*, whereas secretory cells showed increased levels of *MUC5B*, *MUC5AC*, and *SCGB3A1*. Ciliated cells, which are developmentally related to secretory cells, displayed partial expression of secretory markers alongside strong expression of *TMEM190*, *HYDIN*, and *TPPP3*. In addition to canonical epithelial populations, we were able to identify a pathological RAS cell population, characterized by low *SFTPC* and elevated *SCGB3A2* and *SFTPB*. Among stromal populations, adventitial fibroblasts were defined by *PI16*, *MFAP5*, and *SCARA5* expression. CTHRC1+ fibroblasts expressed *CTHRC1*, *POSTN*, and *COL10A1*, while alveolar fibroblasts were marked by *CES1*, *WNT2*, and *TCF21*. Smooth muscle cells (SMCs) and pericytes were identified via *MYH11*, *MYL9*, *CSPG4*, and *RGS5*, respectively. We also distinguished lymphatic endothelial and blood endothelial cells through the expression of *CCL21*, *TFF3*, *MMRN1*, *LYVE1*, *DKK2*, *PLA1A*, and *PLVAP*. Immune populations were robustly represented, including mast cells (*KIT*, *MS4A2*, *SLC18A2*) and CD4⁺ T cells (*CD4*, *CD40LG*, *FOXP3*, *IL7R*). CD8⁺ T cells and natural killer (NK) cells were identified by the expression of *CD8A*, *GZMK*, *NKG7*, *KLRF1*, *KLRD1*, and *GNLY*. Alveolar macrophages and classical monocytes expressed *MERTK*, *CD163*, *S100A9*, and *S100A8*, while *SPP1*+ macrophages co-expressed *SPP1* with tissue-resident markers such as *MARCO*, *C1QA*, and *APOE*. B cells and plasma cells were distinguished by expression of *MS4A1*, *BANK1*, *CD19*, *IGLC1*, *TNFRSF18*, and *JCHAIN*.

### 6. Case study 4: Longitudinal assessment of in-house hPCLS single-cell Flex data

#### 6.1. hPCLS preparation and senescence-induction model

hPCLS were generated as previously described^49^. Briefly, lung tissue cubes (∼1 cm³) were infused via a visible bronchus with warmed 3% UltraPure™ Low Melting Point Agarose (Thermo Fisher, Cat#16520100) prepared in sterile DMEM (Gibco). The infused tissue was transferred to phosphate-buffered saline (PBS) and cooled on ice for 30 min to allow agarose solidification. Embedded tissue was sectioned into 400-µm slices using a vibratome (Leica VT1200; speed 0.10–0.30 mm/s). Slices were incubated at 37 °C with 5% CO₂, washed three times in sterile DMEM/F-12 (Gibco) to remove residual agarose, and cultured overnight in complete DMEM/F-12 supplemented with 10% fetal bovine serum (FBS, Gibco) and 1% Antibiotic-Antimycotic solution (Gibco) prior to experimental use. To establish a bleomycin-induced senescence model, hPCLS from a healthy older donor were randomly assigned to two groups: (1) control (CTRL), cultured in DMEM/F-12 medium, and (2) bleomycin-treated, exposed to 15 µg/mL bleomycin in DMEM/F-12 for 48 hours. After the exposure period, bleomycin was removed, and slices from both groups were maintained in DMEM/F-12 for an additional 72 hours. At the end of the culture period, hPCLS were collected and paraffin-embedded for subsequent scRNA-seq analysis.

#### 6.2 Flex scRNA-seq data generation and analysis

##### 10X Flex scRNA-seq library preparation

Library preparation from bleomycin-treated hPCLS was performed as described in Section 5.1. Briefly, Fixed RNA Profiling was conducted on both untreated and bleomycin-treated hPCLS obtained from a single biological sample from the parenchymal lung region of an aged donor. FFPE blocks were prepared for each sample and preprocessed to assess lung architecture (H&E staining) and RNA integrity (DV200 >30%). Two 50-µm tissue sections were then cut from each FFPE block and placed into 1.5 mL PCR-clean tubes for downstream processing.

##### hPCLS raw sequence mapping and quantification

Raw sequencing reads were processed using the Cell Ranger pipeline (v7.1.0, 10x Genomics) for quality assessment and alignment to the human reference genome. The reference datasets used were refdata-gex-GRCh38-2020-A and Chromium_Human_Transcriptome_Probe_Set_v1.0.1_GRCh38-2020-A provided by 10x Genomics to generate a unique molecular identifier count matrix that was used to create a Seurat object containing a count matrix and analysis. The Seurat object was combined into a merged dataset, and we removed low-quality cells based on high mitochondrial (>10%) and low transcriptional counts (<200). The processed data for each study were imported using R 4.2.3 and Seurat v4.4.0. Samples were integrated using the Seurat and Harmony packages in R, normalized with the SCTransform function using default parameters, scaled, and subjected to dimensionality reduction by principal component analysis on the top 3,000 most variable genes. To correct batch effects and integrate the datasets, the IntegrateData function was applied. For clustering, the FindNeighbors and FindClusters functions were used.

EPCAM^+^ and CDH1^+^ (PTPRC^-^/PDGFRA^-^/PECAM1^-^) were used as canonical gene markers for epithelial cells. Each cell subtype was split into clusters and manually annotated with known cell type markers. *“SFTPC”, “HHIP”, “ABCA3”, “LAMP3”, “PGC”, “SFTPA2”, “SFTPB”, “SERPINA1”* and *“NAPSA”* were used to identify AT2 cells. *“ITGB6”, “ANXA1”, “KRT8”, “KRT18”, “CLDN4”, “SFN”,* and *“KRT7”* were used to identify AT2 transitional cells. *“MMP7”, “CDKN1A”* and *“GDF15”,* were used to identify Aberrant basaloid cells. *“PDPN”, “CAV1”, “CLIC”, “AGER”, “RTKN2”, “HOPX”, “GPRC5A”, “EMP2”, “CLDN18”,* and *“CLIC3”* were used to identify AT1 cells. *“TP63”, “CDH3”, “S100A2”, “KRT5”,* and *“KRT17”* were used to identify Basal cells. *“TPPP3”, “FOXJ1”, “TMEM190”, “CAPS”,* and *“HYDIN”* were used to identify Ciliated cells. *“MUC5B”, “SCGB1A1”, “SCGB3A2”, “SCGB3A1”* and *“BPIFB1”* were used to identify Secretory cells.

PDGFRA^+^ (PTPRC^-^/EPCAM^-^/CDH1^-^/PECAM1^−^) were used to define mesenchymal cells. Each subgroup was then further split into clusters and manually annotated with known fibroblast subtype markers. *“NPNT”, “CES1”, “WNT2”, “TCF21”, “INMT”, “FIBIN”, “AOC3”, “GPM6B”, “SCN7A”, “FMO2”, “CYP4B1”, “LIMCH1”, “FGFR4”, “ITGA8”, “RARRES2”, “GPC3”,* and *“SPINT2”* were used to identify Alveolar fibroblast. *“MFAP5”, “SCARA5”, “PTGIS”, “PI16”, “IGFBP6”,* and *“SFRP2”* were used to identify Adventitial fibroblast. *“CTHRC1”, “TGFBI”, “POSTN”,* and *“COL1A1”,* were used to identify CTHRC1-positive fibroblast. *“CXCL2”* and *“CXCL8”,* were used to identify Inflammatory fibroblast. *“MYH11”, “MYL9”, “ACTA2”,* and *“LMOD1”* were used to identify Smooth muscle cells. *“RGS5”* and *“CSPG4”,* were used to identify Pericytes.

PTPRC^+^ (PDGFRA ^-^/EPCAM^-^/CDH1^-^/PECAM1^−^) were used to define Immune cells. Each subgroup was then further split into clusters and manually annotated with known immune subtype markers. *“SPP1” and “MARCO”* were used to identify SPP1-positive macrophages. *“MSR1”, “C1QA” and “MERTK”* were used to identify Alveolar macrophages. *“S100A9”, “S100A8”, and “CSF3R”,* were used to identify Classical monocytes. *“CD8A”*, *“TRGC2”, “CD8B”, “TNFRSF25”, and “CD40LG”,* were used to identify T cells. *“TRDC”, “NKG7”, “KLRF1”, “KLRD1”,* and *“GNLY”* were used to identify NK cells. *“IGLC1”, “IGHG1”, “JCHAIN”, “TNFRSF18”, “IGKC”, “IGHA1”, “IGHM”,* and *“MZB1”* were used to identify Plasma cells. “KIT”, “CPA3”, and “FCER1A” were used to identify MAST cells. *“MS4A1”, “BANK1”,* and *“CD19”* were used to identify B cells.

PECAM1^+^ (PDGFRA ^-^/EPCAM^-^/CDH1^-^/ PTPRC ^−^) were used to define Endothelial cells. Each subgroup was then further split into clusters and manually annotated with known endothelial subtype markers. *“DKK2”, “GJAS”, “BMX”, and “HEY”* were used to identify EC arterial. *“PROX1” and “CCL21”* were used to identify EC Lymphatic. *“CA4”, “VIPR1”, and “ADGRL2”* were used to identify *aCap cell type*. Doublet cells were identified manually as expressing markers for different cell types, and the final object was created by merging all annotated, doublet-removed subgroups.

### 6.2. Transcription Factor Inference for SnGs in senescence *CTHRC1+* Fibroblast

TF inference was performed for senescence-associated gene sets identified in *CTHRC1+* fibroblasts using CHEA3^50^. *CTHRC1+* fibroblasts-specific senescence-associated genes derived from DeepSAS analysis were used as input. CHEA3 integrates TF–target and co-expression evidence from six independent resources, including ENCODE, ReMap, literature-curated ChIP-seq datasets, ARCHS4 co-expression, GTEx co-expression, and Enrichr libraries. For each TF, CHEA3 assigns a rank within each resource, and an aggregate score was calculated as the mean rank across all six datasets. TFs were subsequently ranked based on this aggregated mean rank for downstream analysis.

## Data Availability

The raw and processed IPF scRNA-seq and Xenium spatial transcriptomics data generated in this study are preserved and will be made available upon publication. Access to the data before publication can be provided upon reasonable request to the corresponding author, Dr. Qin Ma. The human RPE and choroid snRNA-seq atlas is not yet publicly available, as it is currently under review for publication. Access to the data can be provided upon reasonable request to Dr. Rui Chen.

## Code Availability

The DeepSAS source code is available at https://github.com/chthub/deepsas. All codes are publicly available and open source under the MIT License.

## Acknowledgement

This work was supported by NIH grants U54AG075931 (TriState SenNet) and R01GM152585 (QM). Dr. Rui Chen and the generation of the human RPE and choroid snRNA-seq atlas were supported by the Chan Zuckerberg Initiative (CZI) under awards 2019-002425, 2021-239847, and 2021-237885. We thank Dr. Qing Nie (University of California, Irvine) for his invaluable support in helping the cell–cell communication graph design and implementation.

## Author Contributions

*Conceptual design*: Anjun Ma and Qin Ma

*Computational framework design*: Anjun Ma and Qin Ma

*IPF experimental design*: Ana L. Mora and Mauricio Rojas

*IPF sample processing and data analysis*: Natalia Del Pilar Vanegas, Ahmed Ghobashi, Jhonny Rodriguez, and Lorena Rosas

*Choroid sample support and analysis*: Jianming Shao, Ahmed Ghobashi, and Rui Chen,

*Code deployment and optimization*: Hao Cheng and Hu Chen

*Benchmarking and testing*: Yi Jiang, Hu Chen, Xiaoying Wang, and Anjun Ma

*Manuscript draft*: Anjun Ma, Hao Cheng, Ana Mora, Mauricio Rojas, and Qin Ma

*SenNet TriState knowledge support*: Irfan Rahman, Jose Lugo-Martinez, Dongmei Li, Gloria S. Pryhuber, Dongjun Chung, Chi Zhang, Ziv Bar-Joseph, Oliver Eickelberg, Melanie Königshoff, Toren Finkel

All authors revised and approved the final manuscript.

## Ethics declarations

### Competing interests

The authors declare no competing interests.

## Notes

https://github.com/chthub/deepsas

### Competing Interest Statement

The authors have declared no competing interest.

### Summary of Updates

We have extend this work from a short communication to a full research manuscript. This version includes 5 figures. We added new analysis of public healthy lung data and in-house hPLCS data. The biological interpretation focuses on one of the identified CTHRC1+ fibroblast senescence marker, NFE2L2. Added Dr. Chi Zhang as the co-author.

