## Supplementary Information for "Characterizing Cellular Heterogeneity and Transcriptomic Features of Senotype Using Deep Graph Representation Learning"

**Supplementary Tables**

**Supplementary Table S1**: Robustness test results

**Supplementary Table S2**: SnC proportions of benchmarking with other tools

**Supplementary Table S3**: Cell type shared SnGs identified in the human RPE and choroid snRNA-seq data

**Supplementary Table S4**: Cell type specific SnGs identified in the human RPE and choroid snRNA-seq data

**Supplementary Table S5**: Marker gene list used for cell type determination in the human IPF scRNA-seq data

**Supplementary Table S6**: SnC proportions of the human IPF scRNA-seq data

**Supplementary Table S7**: Cell type shared SnGs identified in the human IPF scRNA-seq data

**Supplementary Table S8**: Overlap of shared SnGs with known senescence hallmark lists

**Supplementary Table S9**: Cell type specific SnGs identified in the human IPF scRNA-seq data

**Supplementary Table S10**: Cell type shared SnGs related to age and IPF disease

**Supplementary Table S11**: The gene panel designed for Xenium experiment

**Supplementary Table S12**: CTHRC1+ fibroblast SnG comparison between PCLS and IPF

**Supplementary Table S13**: Pathway enrichment of CTHRC1+ fibroblast SnGs in PCLS and IPF

**Supplementary Table S14**: TF analysis results of CTHRC1+ fibroblast SnGs in IPF and PCLS

**Supplementary Table S15**: Source data information of the human IPF scRNA-seq data

**Supplementary Figures**

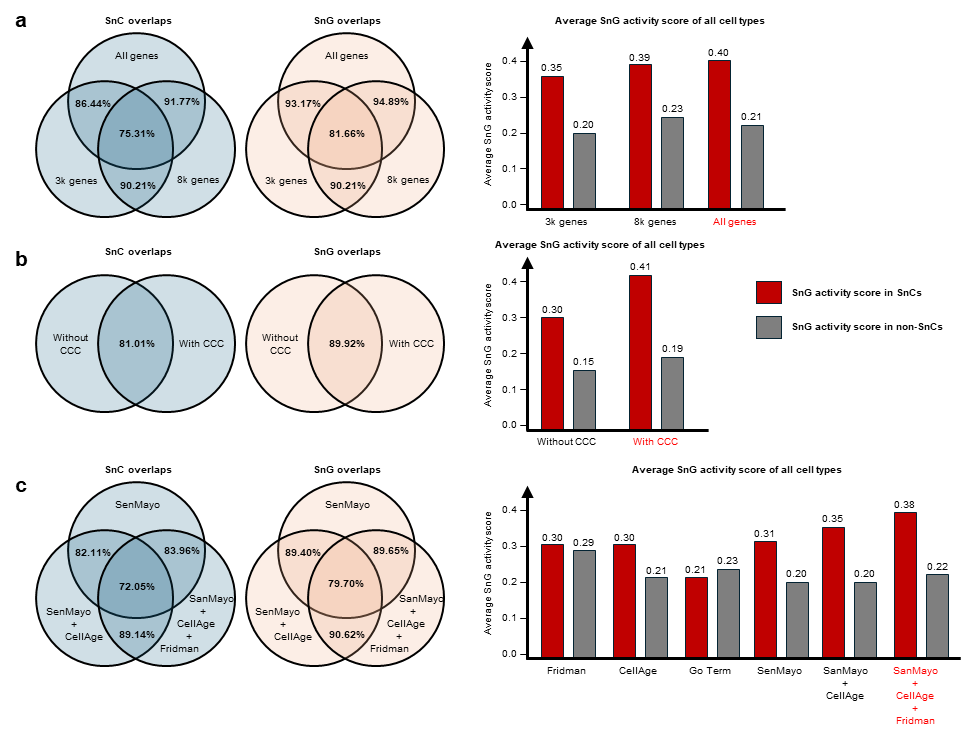

**Supplementary Figure S1**. Results of the ablation test. We evaluated the overlaps of SnCs and SnGs from different ablation tests. SnG activity scores were calculated using Seurat and averaged across all cell types. The higher the activity, the better. Words in red indicate default settings used in DeepSAS. (a) the number of genes used to construct the heterogeneous graph, (b) the inclusion of CCC information, and (c) the use of different hallmark gene lists for labeling initial SnG candidates.

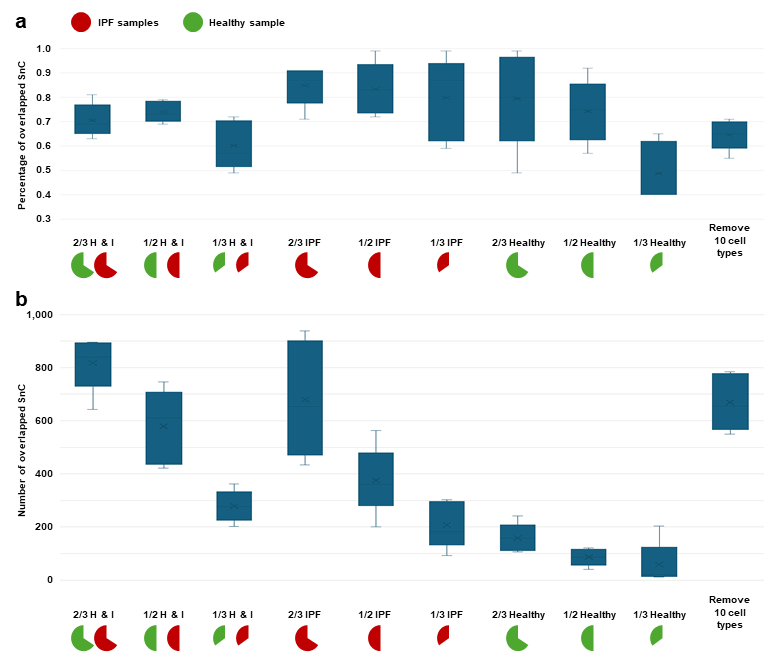

**Supplementary Figure S2**. Robustness test results. SnCs predicted from subsampling batches were evaluated by comparing with the original full data results. Each box represents the results of five replicate experiments. (a) percentage of overlapping SnCs. (b) Number of overlapping SnCs.

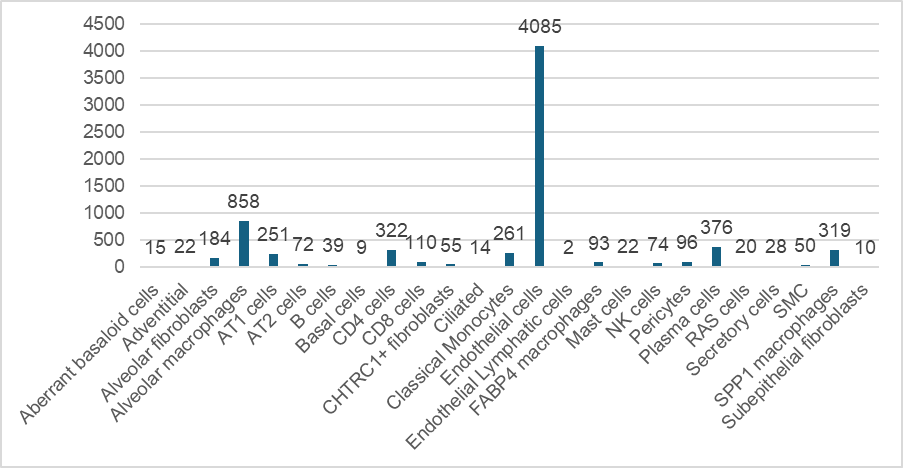

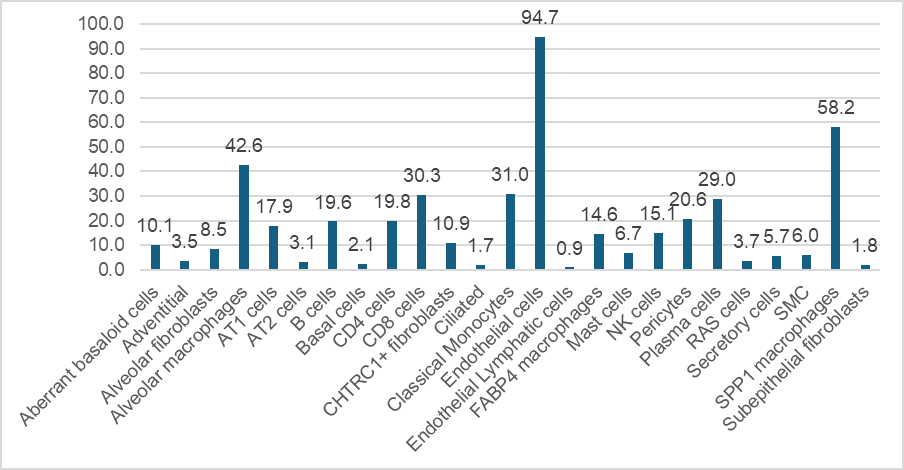

**DeepSAS**

**MarsGT+ChatGPT**

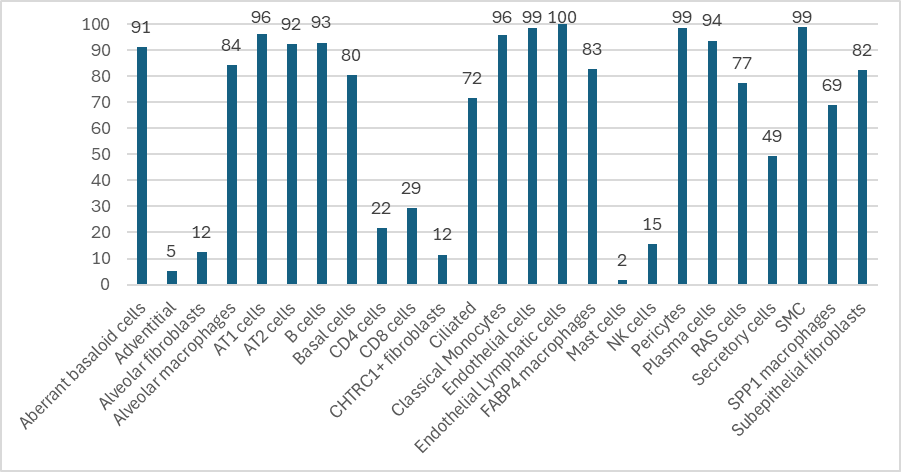

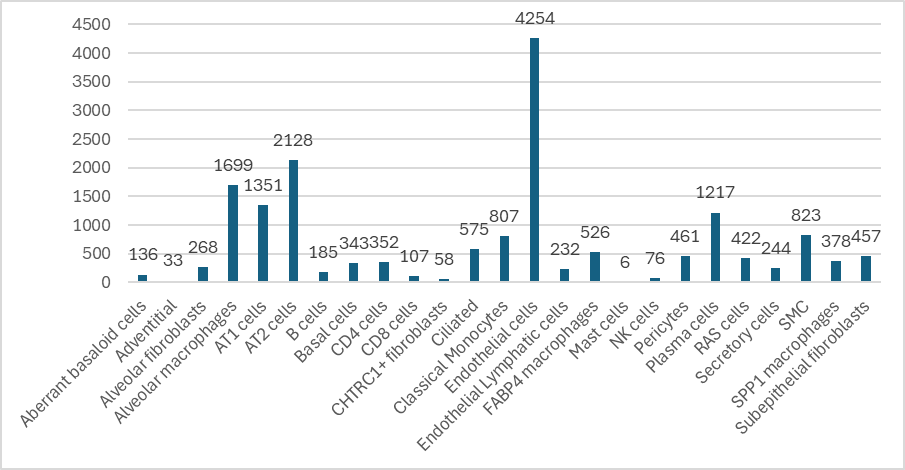

**Seurat+ChatGPT**

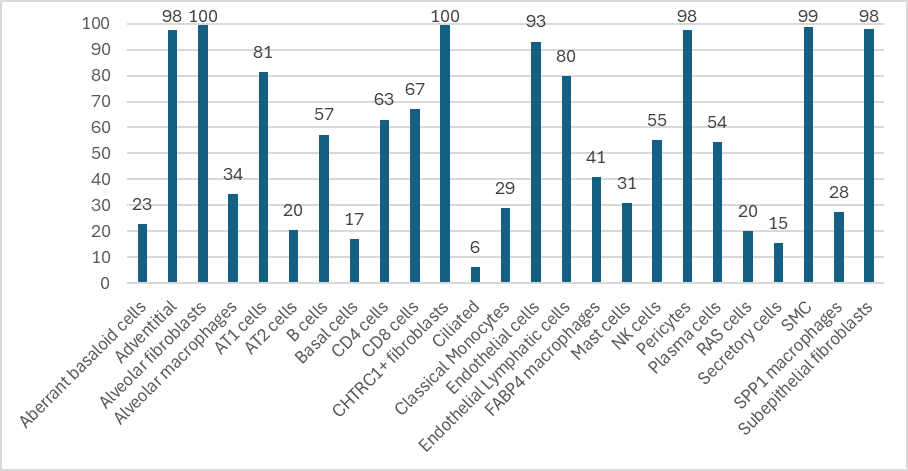

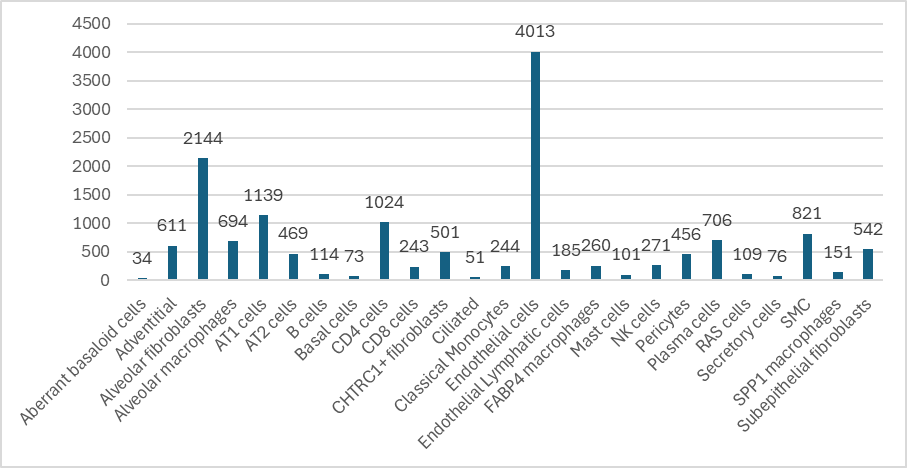

**SenCID**

**SenePy**

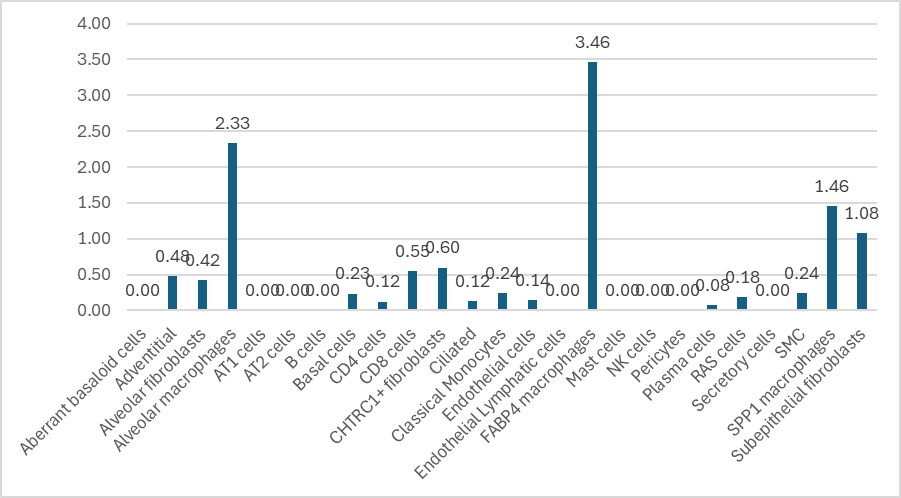

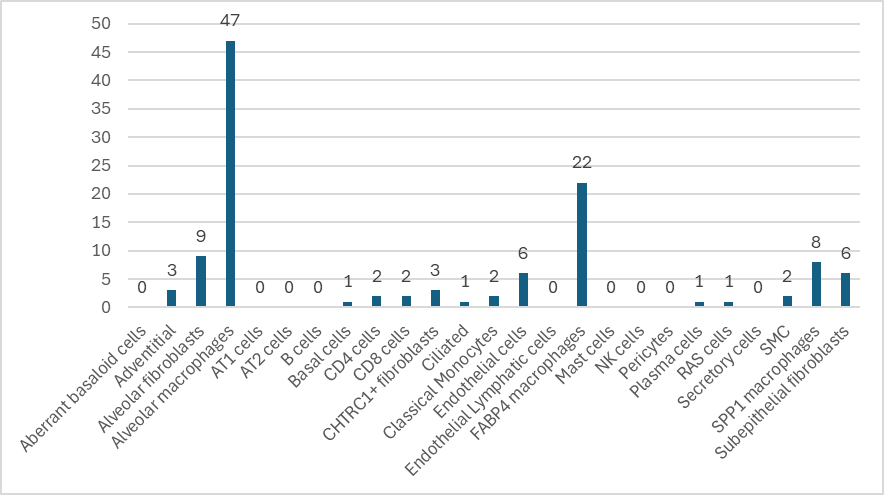

**Number of SnCs**

**Percentage of SnCs**

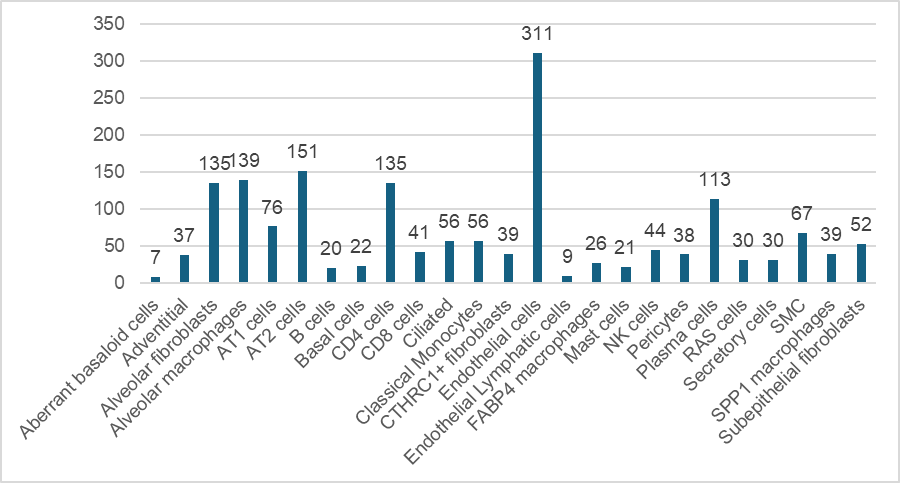

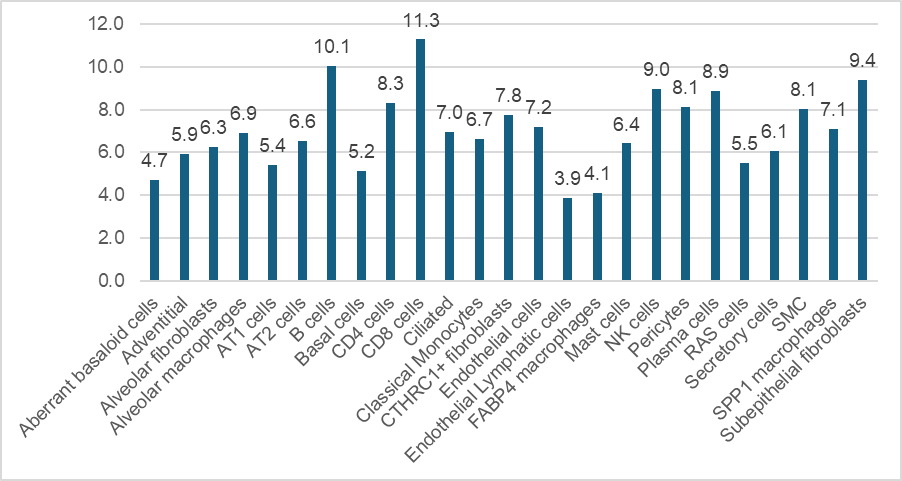

**Supplementary Fig. S3.** Comparison of SnC prediction results of different tools. The number of SnCs and the proportion of SnCs were showcased, divided by cell types.

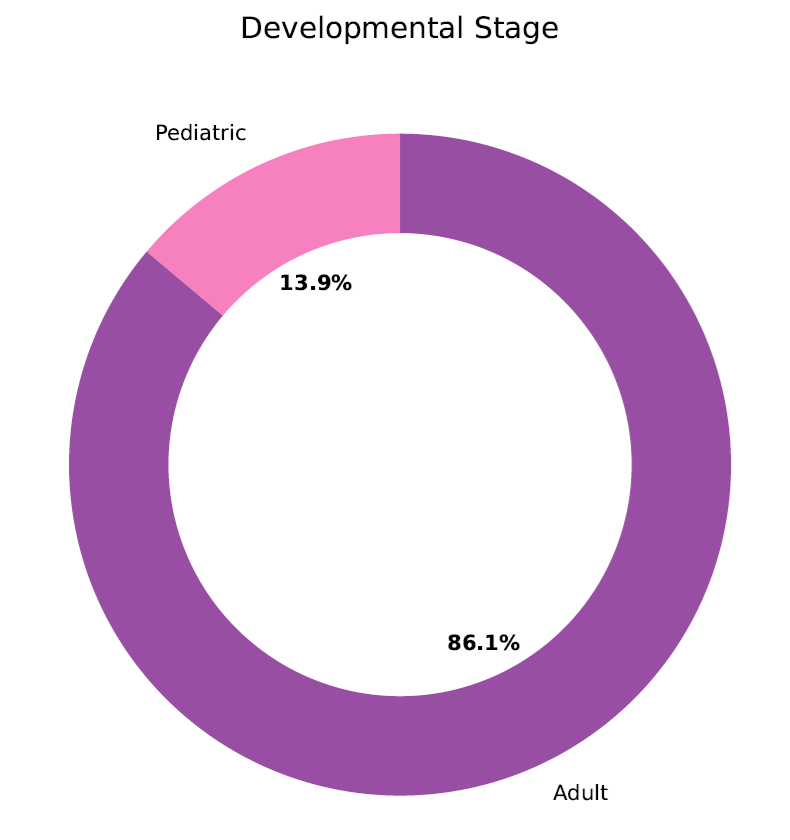

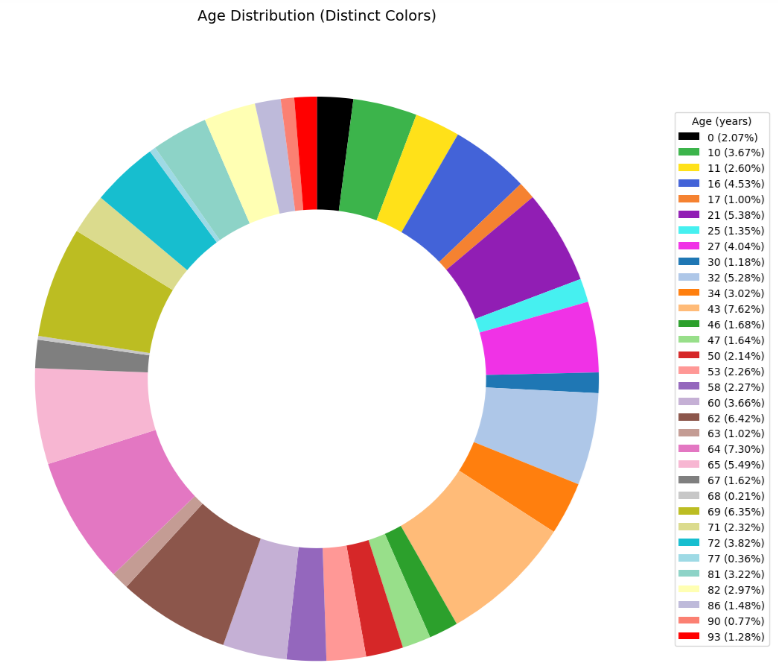

**a**

**b**

**Supplementary Fig. S4.** Percentage of developmental stage and ages of the RPE and choroid single-cell atlas.

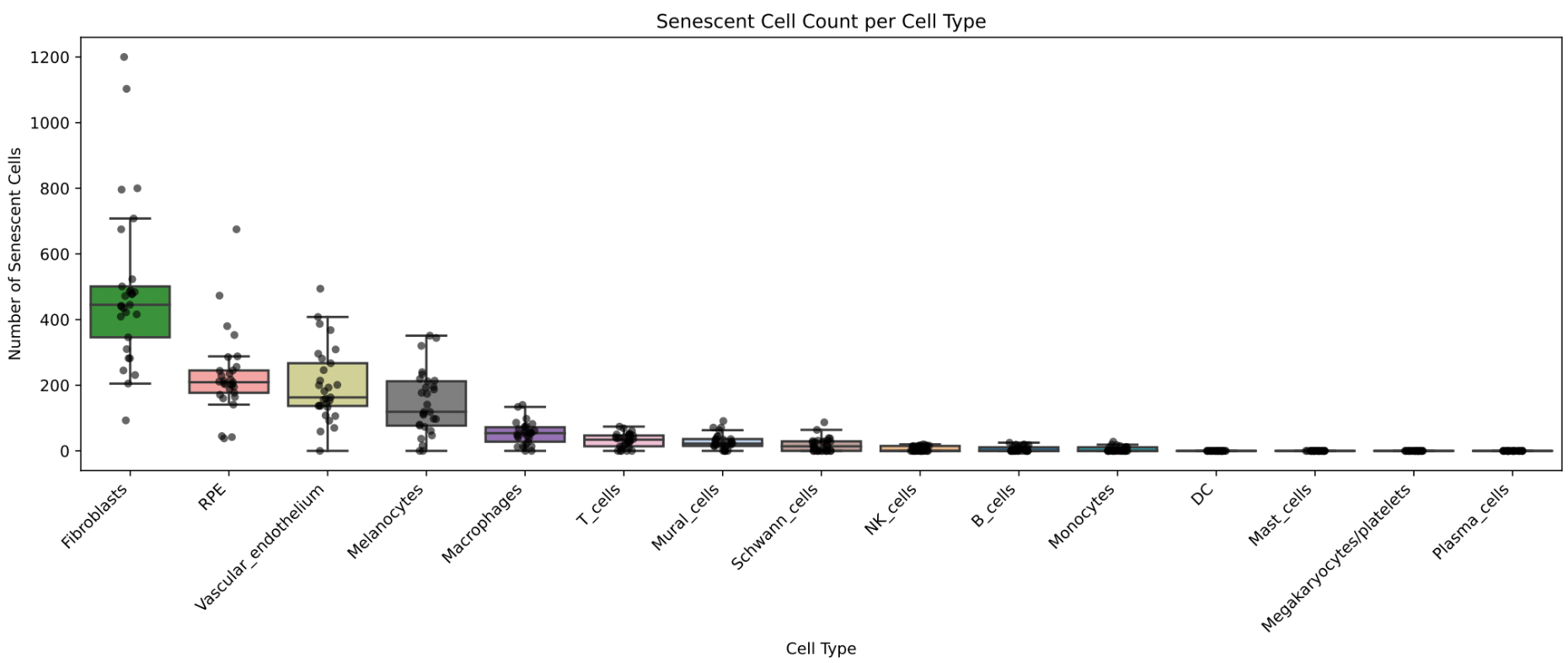

**Supplementary Fig. S5.** Number of SnCs across 15 cell types in the RPE and choroid single-cell atlas, colored by age group. Each dot represents a batch.

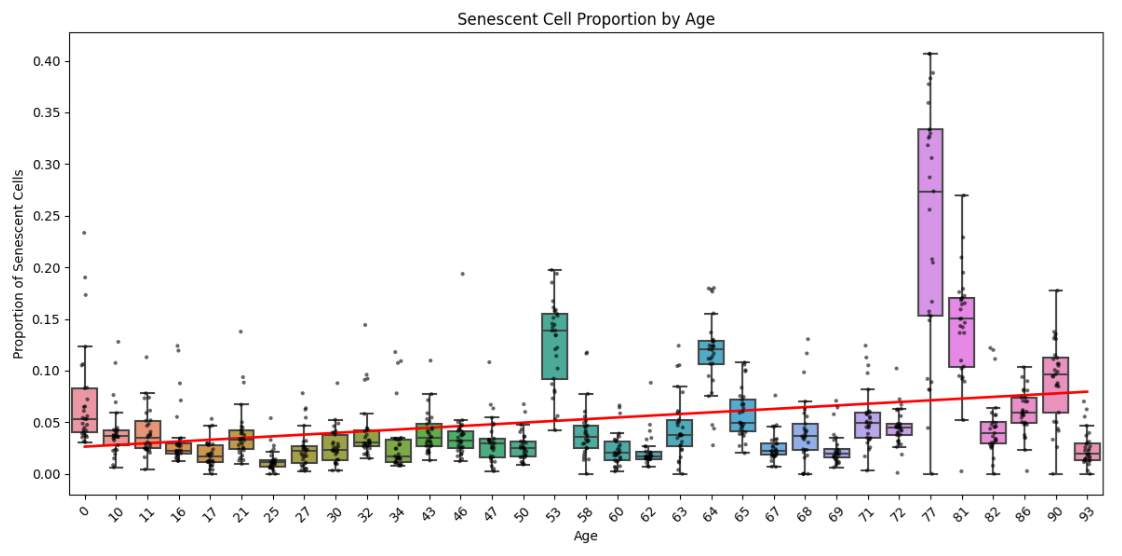

**Supplementary Fig. S6.** Boxplots comparing DeepSAS predicted SnC proportions across age groups, combining results in all cell types. Each dot represents a batch.

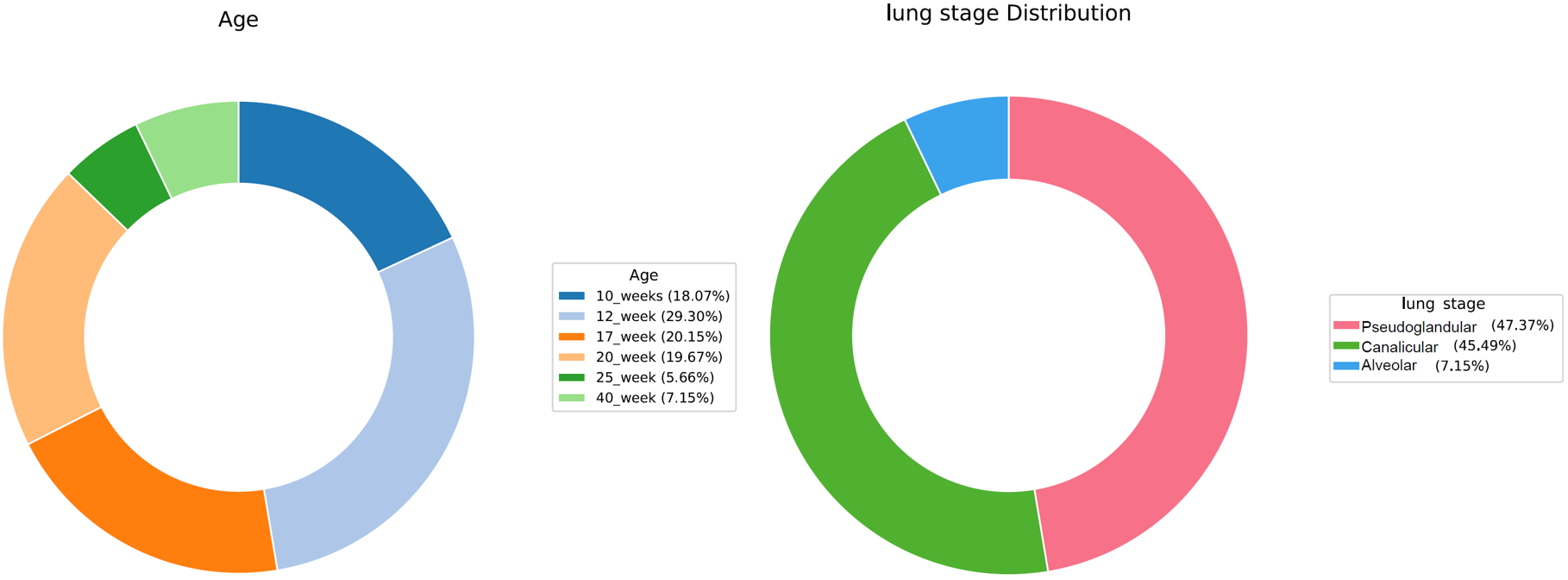

**Supplementary Fig. S7.** Percentage of sample distribution of public healthy lung data.

Bronchiolo-vascular bundles

Dense hyalinized fibrous scarring

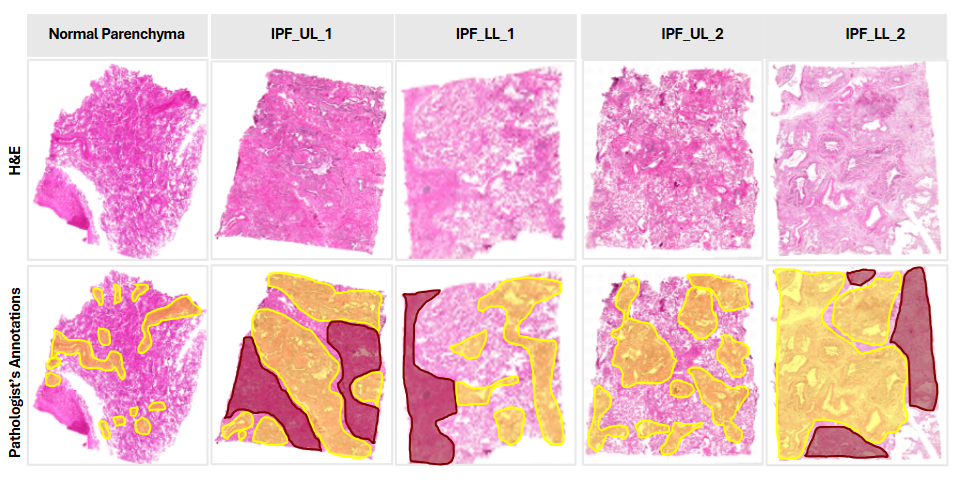

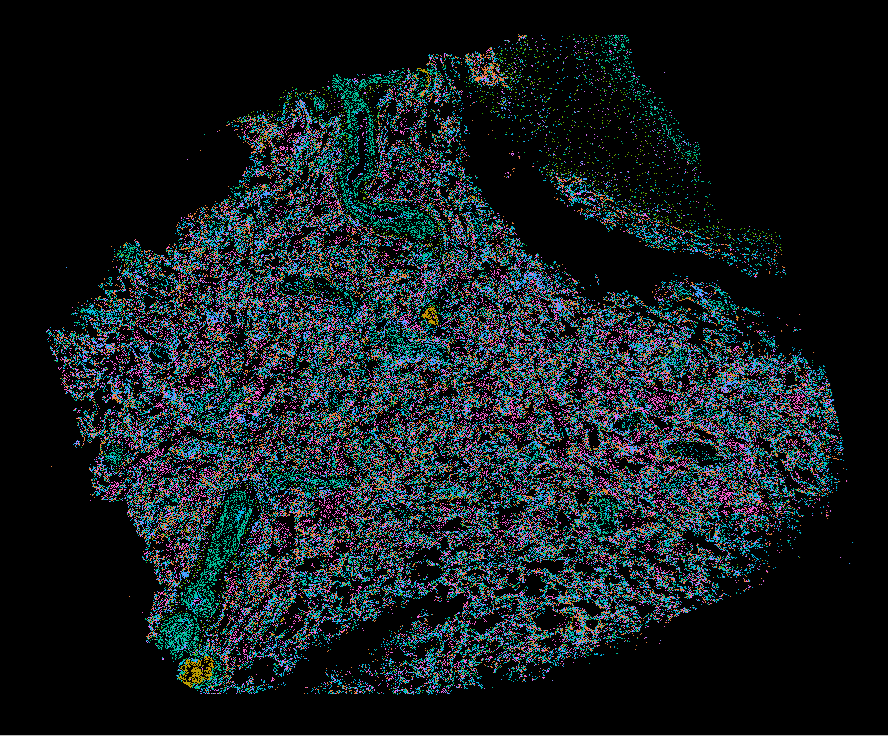

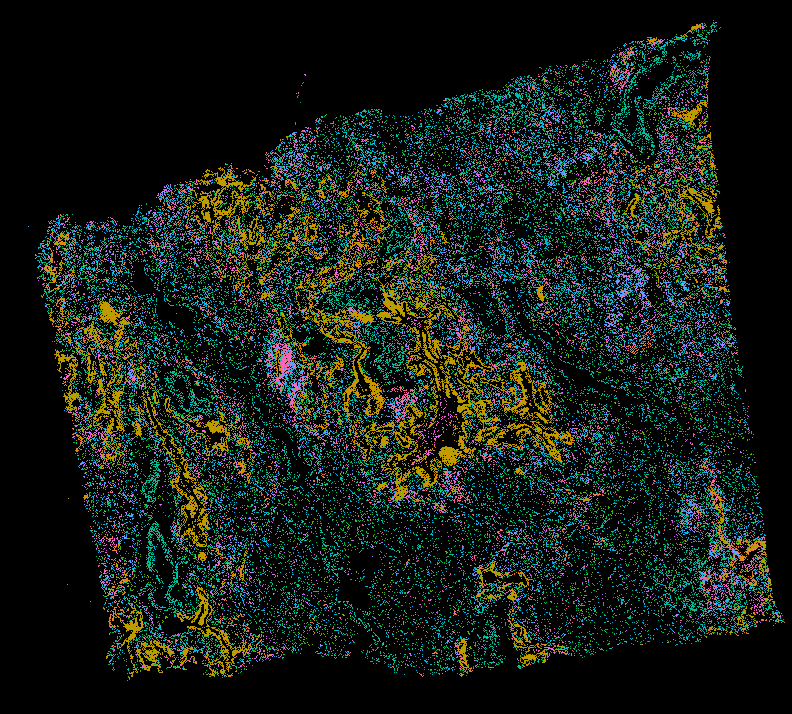

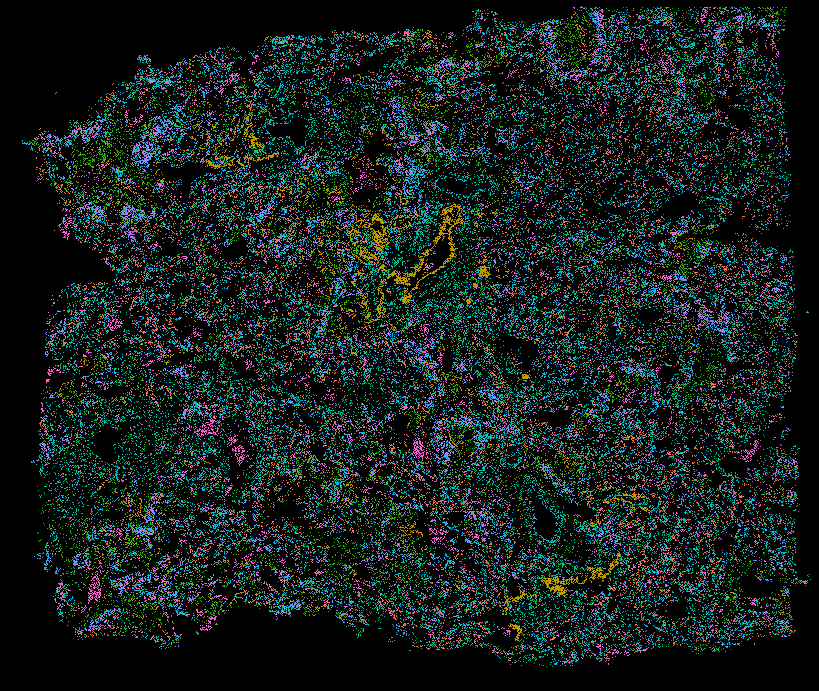

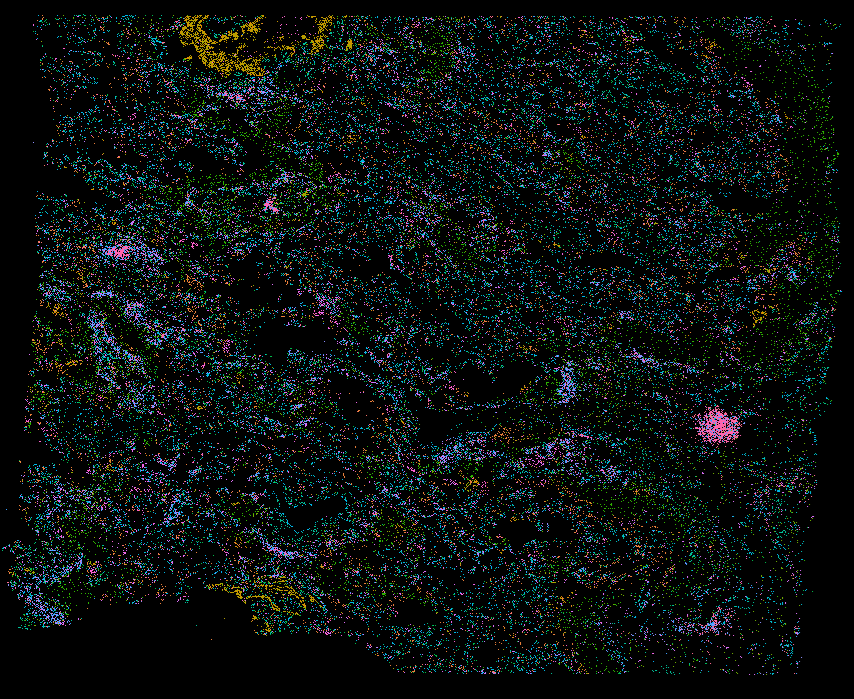

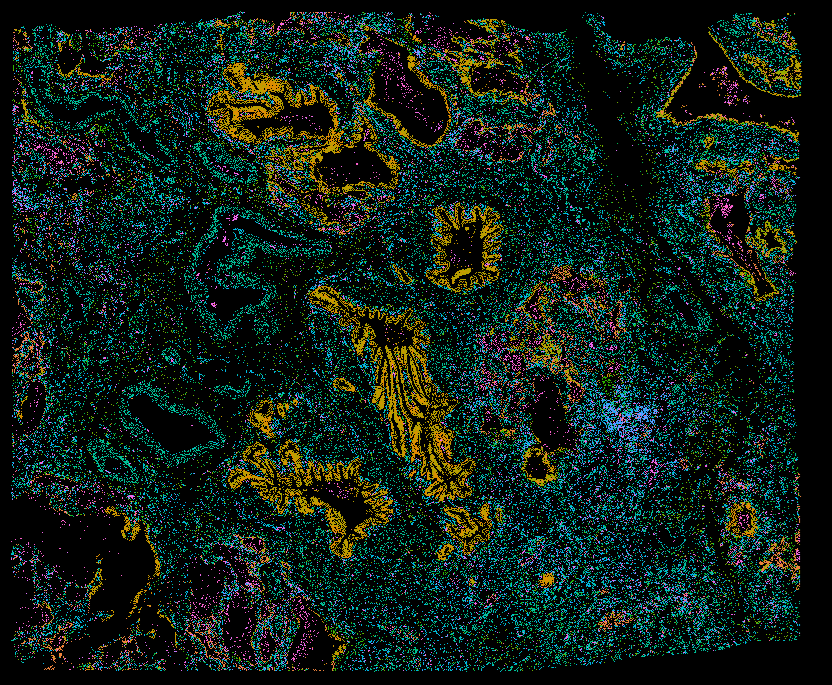

**H&E**

**Pathological annotation**

**Panel gene expression**

**Normal parenchyma**

**IPF_UL_1**

**IPF_LL_1**

**IPF_UL_2**

**IPF_LL_2**

**Supplementary Fig. S8.** The whole Xenium slides of all five samples are provided. The original H&E image, the image with pathological annotation, and the image colored by panel gene expressions are showcased.

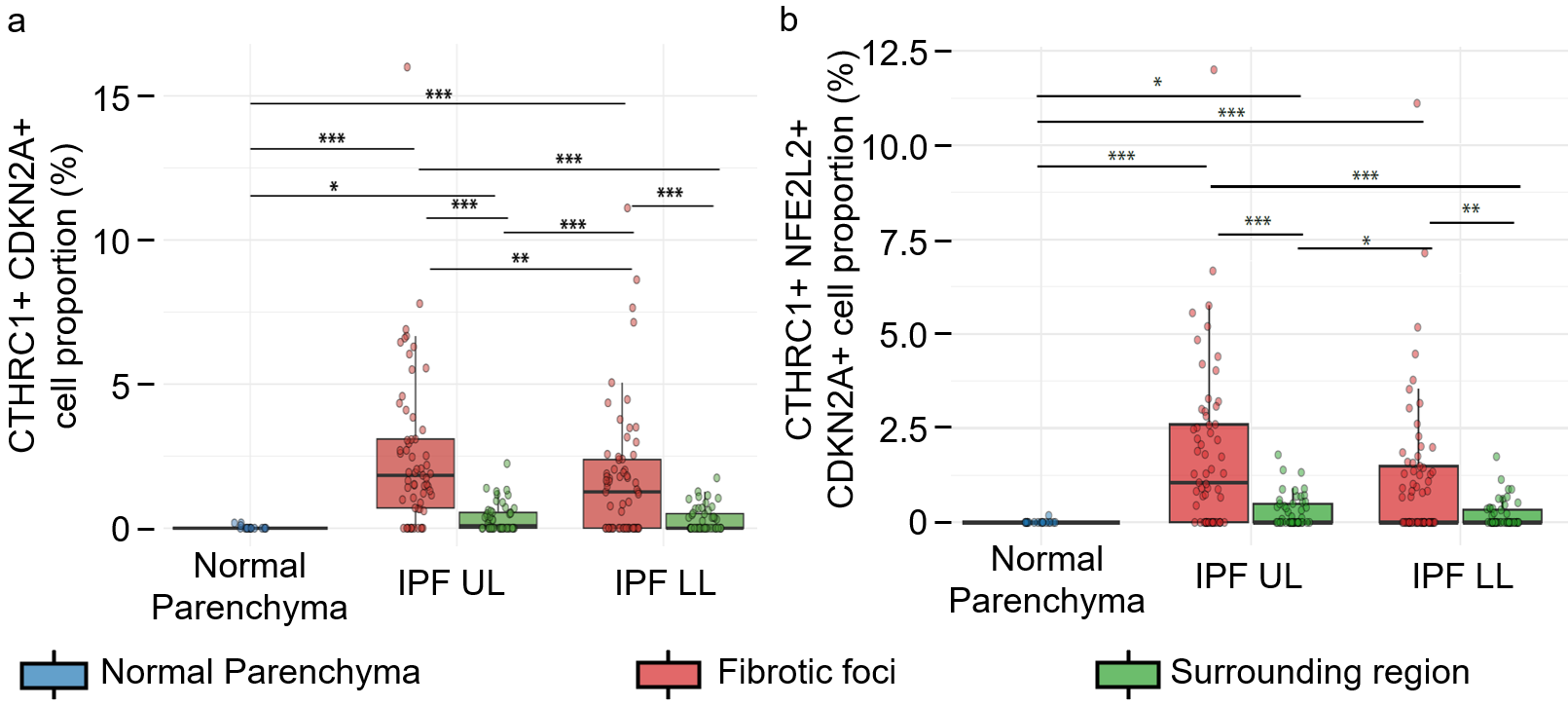

**Supplementary Fig. S9.** Quantification of co-expression of *CTHRC1* and *CDKN2A* **(a)** and *CTHRC1*, *NFE2L2*, and *CDKN2A* **(b)** cell proportions across selected regions in normal parenchyma, IPF upper lobe, and IPF lower lobe samples. Box plots summarize results from 19 normal regions, 126 fibrotic foci regions, and 101 surrounding regions. Statistical significance was assessed using the Kruskal–Wallis test. *p* < 0.05 (*), *p* < 0.01 (**), and *p* < 0.001 (***)

**Supplementary Fig. S10.** Xenium validation of IL6 and CCL4 expressions. (a) An example of the co-expression of IL6 and CCL4 in the surrounding regions. (b) Boxplot showing significantly increased abundance of double-positive (CCL4 and IL6) cells in surrounding regions relative to fibrotic foci in control and both lobes. Statistical significance was assessed using the Kruskal–Wallis test. *p* < 0.05 (*), *p* < 0.01 (**), and *p* < 0.001 (***)

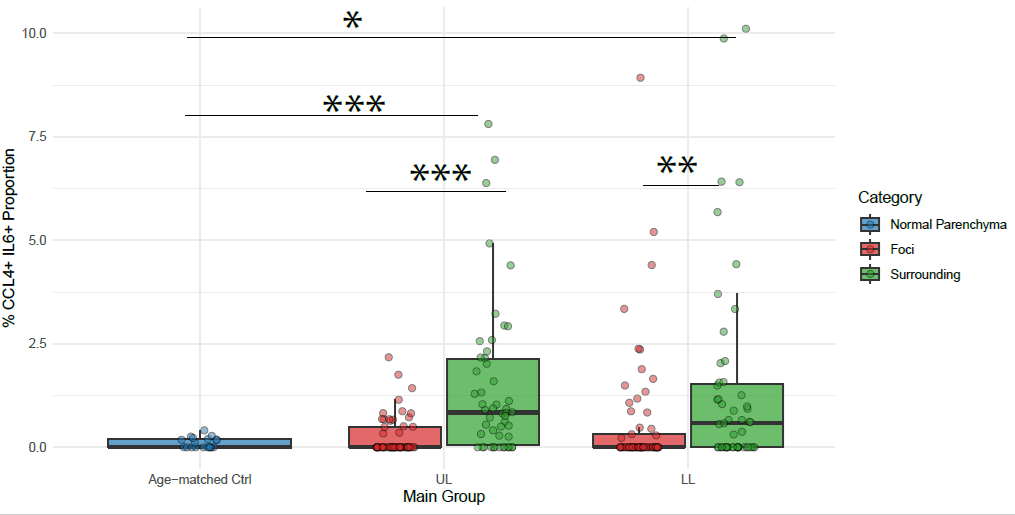

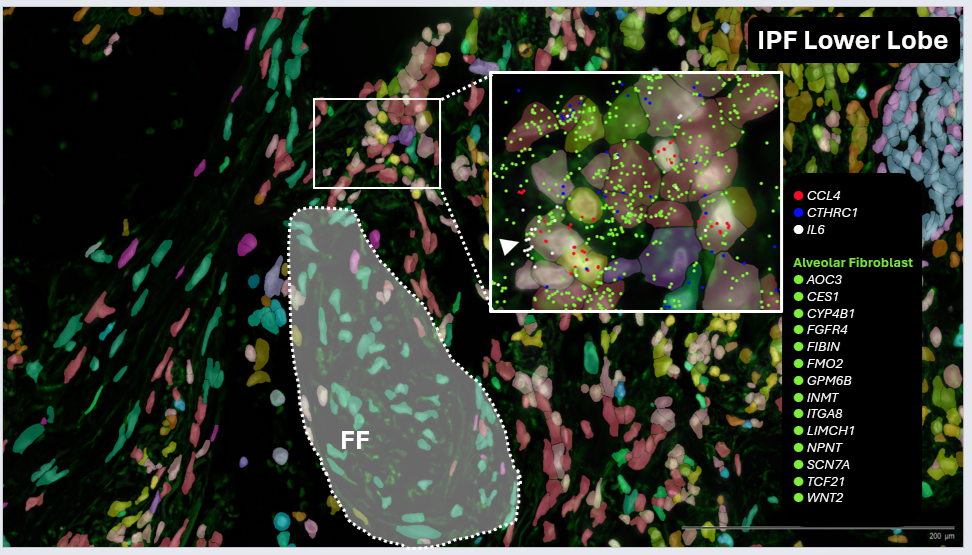

**a**

**b**

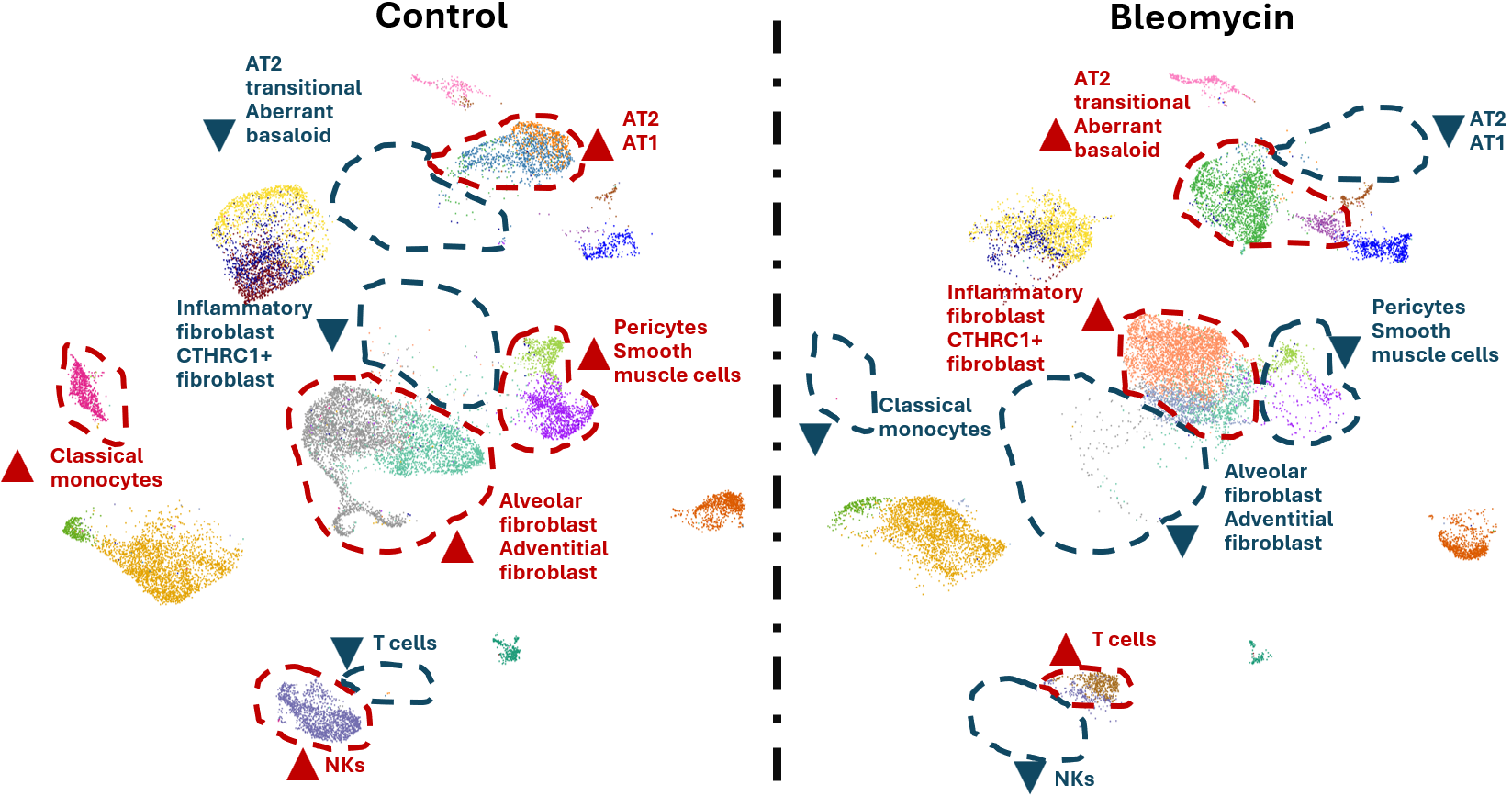

**Supplementary Fig. S11.** UMAP of the PCLS scRNA-seq data, split by the control and bleomycin-treated samples. Red triangle on one side indicates an increase in cell number in the specific cell types, and blue triangle means a decrease in cell numbers in the cell type, compared to the other sample.
